# Genetic risk underlying psychiatric and cognitive symptoms in Huntington’s Disease

**DOI:** 10.1101/639658

**Authors:** Natalie Ellis, Amelia Tee, Branduff McAllister, Thomas Massey, Duncan McLauchlan, Timothy Stone, Kevin Correia, Jacob Loupe, Kyung-Hee Kim, Douglas Barker, Eun Pyo Hong, Michael J. Chao, Jeffrey D. Long, Diane Lucente, Jean Paul G. Vonsattel, Ricardo Mouro Pinto, Kawther Abu Elneel, Eliana Marisa Ramos, Jayalakshmi Srinidhi Mysore, Tammy Gillis, Vanessa C. Wheeler, Christopher Medway, Lynsey Hall, Seung Kwak, Cristina Sampaio, Marc Ciosi, Alastair Maxwell, Afroditi Chatzi, Darren G. Monckton, Michael Orth, G. Bernhard Landwehrmeyer, Jane S. Paulsen, Ira Shoulson, Richard H. Myers, Erik van Duijn, Hugh Rickards, Marcy E. MacDonald, Jong-min Lee, James F. Gusella, Lesley Jones, Peter Holmans

## Abstract

Huntington’s disease (HD) is an inherited neurodegenerative disorder caused by an expanded CAG repeat in the *HTT* gene. It is diagnosed following a standardized exam of motor control and often presents with cognitive decline and psychiatric symptoms. Recent studies have detected genetic loci modifying the age at onset of motor symptoms in HD, but genetic factors influencing cognitive and psychiatric presentations are unknown. We tested the hypothesis that psychiatric and cognitive symptoms in HD are influenced by the same common genetic variation as in the general population by constructing polygenic risk scores from large genome-wide association studies of psychiatric and neurodegenerative disorders, and of intelligence, and testing for correlation with the presence of psychiatric and cognitive symptoms in a large sample (n=5160) of HD patients. Polygenic risk score for major depression was associated specifically with increased risk of depression in HD, as was schizophrenia risk score with psychosis and irritability. Cognitive impairment and apathy were associated with reduced polygenic risk score for intelligence. In general, polygenic risk scores for psychiatric disorders, particularly depression and schizophrenia, are associated with increased risk of the corresponding psychiatric symptoms in HD, suggesting a common genetic liability. However, the genetic liability to cognitive impairment and apathy appears to be distinct from other psychiatric symptoms in HD. No associations were observed between HD symptoms and risk scores for other neurodegenerative disorders. These data provide a rationale for treatments effective in depression and schizophrenia to be used to treat depression and psychotic symptoms in HD.

## Introduction

Huntington’s disease (HD) is an inherited neurodegenerative disorder caused by an expanded CAG repeat in *HTT*. A clinical diagnosis is typically made via a movement disorder, but nearly all participants experience progressive cognitive decline, and many exhibit behavioural and psychiatric symptoms (1). Depression, irritability, obsessive and compulsive symptoms, apathy and psychosis all occur at rates higher than seen in the non-HD population, though they are not universal in HD (2). Psychiatric symptoms are often present before motor symptoms become manifest. Age at motor onset of HD is determined both by the length of the CAG repeat tract in *HTT* and by other genetic variants in the genome (3–5). Despite the fact that age at motor onset measures only one specific facet of the pathological process (1), it has been widely used to identify genetic modifiers in HD whilst genetic influences on behavioural and neuropsychiatric symptoms in HD have not been systematically investigated. Small studies have shown familial aggregation of psychosis in HD (6, 7) with weak evidence for the influence of specific candidate genes (8).

Common genetic variation contributes to the risk of developing schizophrenia (SCZ) (9), bipolar disorder (BPD) (10), major depressive disorder (MDD) (11) and attention deficit hyperactivity disorder (ADHD) (12). There is significant shared genetic risk between these psychiatric disorders (13).

Increased general intelligence (*g*) – a measure of cognitive function - has been shown to be genetically correlated with reduced risk of Alzheimer’s disease (AD), ADHD and schizophrenia (14). As in HD, there are substantially increased levels of psychiatric symptoms in many neurological diseases. For instance 50% of those with AD (15), and up to 75% of those with Parkinson’s disease (PD) develop psychotic symptoms (16, 17). In dementia with Lewy bodies visual hallucinations are a core clinical feature seen in 80% of patients (18). AD with psychosis is heritable (19, 20), though increasing polygenic risk score for schizophrenia was associated with reduced risk of psychosis in PD and AD (21).

Given the increased frequency of neuropsychiatric and cognitive symptoms in HD, it is of interest to test for genetic overlap of these symptoms with psychiatric and neurodegenerative disorders and intelligence. This was done by constructing polygenic risk scores using the latest available genome-wide association studies for these disorders and testing these for correlation with the presence of neuropsychiatric and cognitive symptoms in HD.

## Materials & Methods

### HD participants and phenotypes

The HD participants in this analysis were part of the European REGISTRY study (22) or its successor, the international Enroll-HD (23) observational study of HD. REGISTRY was a multisite, prospective, observational study, which collected phenotypic data (2003–13) for more than 13000 participants, mostly HD gene carriers with manifest disease. Enroll-HD is an expanded and modified version of the REGISTRY study and is international: to date it has over 16000 participants (some of whom rolled over from REGISTRY) from 19 nations. All experiments were performed in accordance with the Declaration of Helsinki, ethical approval for the REGISTRY and Enroll-HD studies including written informed consent of all participants was obtained. This study was approved by Cardiff University School of Medicine Research Ethics Committee.

There were 6278 individuals with manifest HD defined by motor onset from REGISTRY (n=4986) and Enroll-HD (n=1292) with appropriate quality-controlled genome-wide association study (GWAS) data as described in (5). We examined seven symptoms: depression, irritability, psychosis, apathy, violent/aggressive behaviour, perseverative/obsessive behaviour and cognitive impairment, from the clinical characteristics questionnaire (CCQ) in REGISTRY and Enroll-HD. The clinical characteristics questionnaire asks if a sign or symptom has ever been experienced by a subject, (see Supplementary Information Appendix A). At least one CCQ symptom endorsement (positive or negative) was recorded in 5854 participants (4563 REGISTRY, 1291 Enroll-HD). We removed 133 individuals with a comorbid diagnosis of bipolar disorder, schizophrenia, schizotypy or schizoaffective disorder (since these are likely to share risk genes for psychiatric disorders independently of their HD status). We also removed one member (561 individuals) of each pair of first or second degree relatives (IBD>0.25) to minimise the correlation between individuals due to cryptic relatedness. This left 5160 participants in the final analysis (**Supplementary Figure 1**).

**Figure 1.**
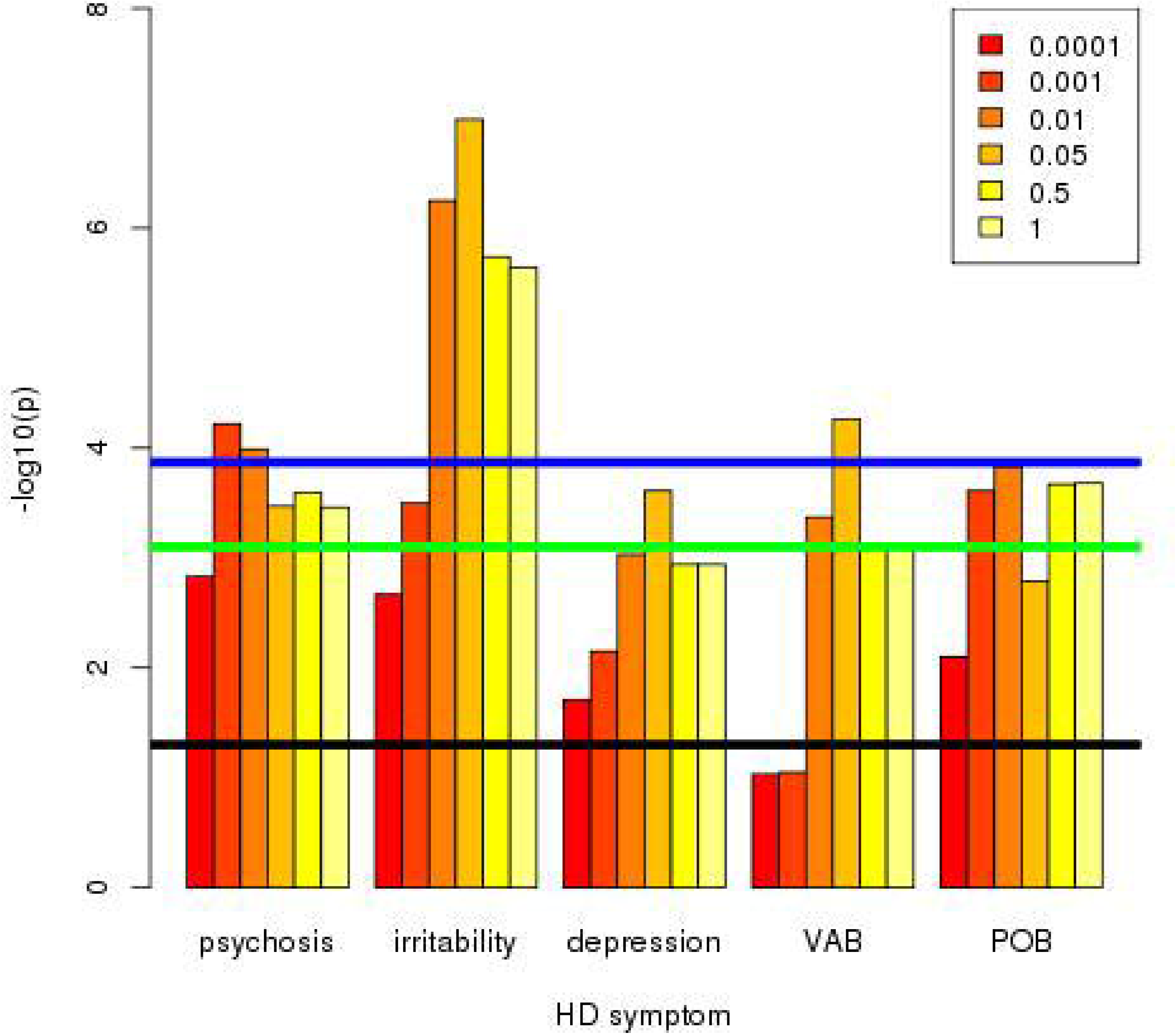
Association of increased schizophrenia PRS with increased HD symptom frequency. Numbers in the box correspond to the p-value thresholds used to derive the PRS. Black line corresponds to nominally significant association (p=0.05). Green line indicates associations significant after Bonferroni correction for 63 PRS-symptom comparisons (9 PRS x 7 symptoms). Blue line indicates associations significant after Bonferroni correction for 63 PRS-symptom comparisons and 6 PRS cutoffs. Note: Only symptoms with at least one p-value reaching the green line are shown.

### Genetic analysis

For each individual in the HD dataset, their genetic risk for each psychiatric/neurodegenerative disorder was captured by a polygenic risk score (PRS) (24). A PRS is defined as the sum of the number of minor alleles across a set of SNPs, each weighted by the corresponding risk of that allele for the psychiatric/neurodegenerative disorder observed in a “training” GWAS. The set of SNPs used to calculate the PRS was chosen to be present in both the training GWAS and the HD dataset, to be in approximate linkage equilibrium, and to capture as much of the association signal in the training GWAS as possible. Following the procedure outlined by the Psychiatric Genomics Consortium (9), this was achieved by taking the most significant SNP in the training GWAS, removing all SNPs within 500kb that are in linkage disequilibrium (r^2^>0.1) with it, then moving to the next most significant remaining SNP and repeating the process. Analysis was restricted to SNPs with minor allele frequency >0.01 that were well imputed (r^2^ between allele dosages and the unknown true genotypes >0.9) in the HD dataset and (where this information was available) in the training GWAS. SNPs were selected for inclusion into the PRS by applying criteria to their p-values in the training set. Since the optimal criterion was not known *a priori*, we applied six different p-value cutoffs (p<0.0001, p<0.001, p<0.01, p<0.05, p<0.5, p<1). Effects of population stratification were removed by regressing the PRS on 20 principal components. The residuals from this regression were standardised, to enable comparison of effect sizes across cutoffs.

Nine large publicly available sets of GWAS summary statistics were used for training (**Table 1**). These comprised five GWAS of psychiatric disorders from the Psychiatric Genomics Consortium (PGC): schizophrenia, bipolar disorder (BPD), attention deficit hyperactivity disorder (ADHD), autism spectrum disorder (ASD) and obsessive compulsive disorder (OCD) (9, 10, 12, 25, 26), a meta-analysis of the PGC and UK Biobank major depressive disorder (MDD) samples (11), two neurodegenerative disorders: late-onset Alzheimer’s disease (AD) (27), and Parkinson’s disease (PD) (28, 29), and a large GWAS of *g*, a measure of general intelligence (14). The major histocompatibility complex (MHC) region was removed from the schizophrenia GWAS due to a strong schizophrenia signal and long-range linkage disequilibrium potentially biasing the PRS (9). Summary statistics for the Psychiatric Genomics Consortium GWAS are available from https://www.med.unc.edu/pgc/results-and-downloads, those from the PGC and UK Biobank MDD meta-analysis from https://datashare.is.ed.ac.uk/handle/10283/3203, those for the AD and intelligence GWAS from https://www.ebi.ac.uk/gwas/downloads/summary-statistics, and the PD GWAS (omitting 23 and Me samples) from https://drive.google.com/drive/folders/10bGj6HfAXgl-JslpI9ZJIL_JIgZyktxn.

**Table 1.** Sample sizes of GWAS used as training sets for polygenic risk scores.

| **<u>DISORDER</u>** | **<u>NUMBER OF CASES</u>** | **<u>NUMBER OF CONTROLS</u>** |
| --- | --- | --- |
| Major depressive disorder (MDD) | 170,756 | 329,443 |
| Bipolar disorder (BPD) | 20,352 | 31,358 |
| Schizophrenia (SCZ) | 36,989 | 113,075 |
| Attention deficit hyperactivity disorder (ADHD) | 20,183 | 35,191 |
| Autism spectrum disorder (ASD) | 18,382 | 27,969 |
| Obsessive-compulsive disorder (OCD) | 2,688 | 7,037 |
| Alzheimer's Disease (AD) | 17,008 | 37,154 |
| Parkinson's Disease (PD) | 26,421 | 442,271 |
| Intelligence | 269,867 individuals |  |

Associations were tested between all 9 PRS and all 7 symptoms at each PRS cutoff. The following p-value criteria were used to define significance correcting for multiple testing: 7.94×10^-4^ (Bonferroni corrected for 7 symptoms tested on 9 disease PRS, a total of 63 tests), 1.32×10^-4^ (Bonferroni corrected for 63 tests x 6 PRS cutoffs)

We also defined 27 PRS-symptom comparisons as primary hypotheses of interest, reflecting prior associations in the general population between the phenotypes and the diseases from which the PRS were derived. These were:

- MDD with depression, irritability and apathy
- BPD with depression and irritability
- Schizophrenia with depression, irritability, psychosis, apathy and violent/aggressive behaviour
- ADHD with irritability, violent & aggressive behaviour and cognitive impairment
- ASD with irritability, psychosis and perseverative/obsessive behaviour
- OCD with perseverative/obsessive behaviour
- AD with depression, irritability, apathy, violent/aggressive behaviour and cognitive impairment
- PD with depression, irritability, apathy and cognitive impairment
- Intelligence with cognitive impairment

## Results

The frequency of each symptom in the 5854 HD participants with at least one endorsed symptom (positive or negative) varied from 10.8% (psychosis) to 66.0% (depression), and differed significantly by sex for depression, which was more common in women, and irritability and violent/aggressive behaviour which were both more common in men (**Table 2**). Symptoms were significantly (positively) correlated with each other (**Supplementary Table 1**), resulting in individuals exhibiting multiple symptoms.

**Table 2:** HD symptom counts and sex differences in the 5854 individuals with at least one recorded HD symptom diagnosis (positive or negative).

| Symptom | Freq (%) | Male |  | Female |  | OR(F/M) | P(F/M) |
| --- | --- | --- | --- | --- | --- | --- | --- |
|  |  | + | - | + | - |  |  |
| Depression | 66.0 | 1639 | 1120 | 2116 | 816 | 1.75<br>(1.57,1.96) | <2e-16 |
| Irritability | 60.3 | 1757 | 1005 | 1675 | 1252 | 0.77<br>(0.69,0.85) | 8.76e-7 |
| Psychosis | 10.8 | 299 | 2396 | 301 | 2562 | 0.94<br>(0.79,1.12) | 0.485 |
| Violent/aggressive behaviour (VAB) | 31.2 | 965 | 1794 | 802 | 2107 | 0.71<br>(0.63,0.79) | 1.90e-9 |
| Apathy | 53.8 | 1484 | 1291 | 1581 | 1344 | 1.02<br>(0.92,1.14) | 0.664 |
| Perseverative/obsessive Behaviour (POB) | 37.5 | 1048 | 1713 | 1076 | 1833 | 0.96<br>(0.86,1.07) | 0.451 |
| Cognitive impairment | 57.5 | 1583 | 1214 | 1726 | 1235 | 1.07<br>(0.97,1.19) | 0.194 |

Freq(%) is the frequency of the symptom among individuals in the sample for whom a diagnosis of that symptom was recorded. +/- refer to the number of individuals of each sex with/without the symptom.

We approximated disease duration as the age at the most recent observation available minus age at motor onset for REGISTRY and Enroll-HD individuals. Mean duration of HD was significantly lower in Enroll-HD, at 6.54 years, than REGISTRY, at 8.14 years (p=6.55×10^-21^). The effects of CAG length, age at motor onset, sex and disease duration on symptom presence, were tested simultaneously via logistic regression (**Table 3**), along with potential differences in symptom frequency between REGISTRY vs Enroll-HD. Increased disease duration was significantly correlated with symptom presence for all symptoms except depression. These associations remained significant when age at most recent visit was included in the model (to account for increasing frequency of symptoms with increasing age) instead of age at motor onset (results not shown). Thus, the increased frequency of symptoms with disease duration cannot simply be attributed to age. Perseverative/obsessive behaviour was seen substantially more frequently in Enroll-HD than REGISTRY participants (p = 5.72×10^-10^), otherwise there were no significant differences.

**Table 3:** Effect of sex, CAG, age at motor onset, disease duration and sample collection (ENROLL vs REGISTRY) on HD symptom frequency

| HD Symptom | Sex (F/M) |  | CAG |  | AMO |  | Duration |  | ENR vs REG |  | AUC |
| --- | --- | --- | --- | --- | --- | --- | --- | --- | --- | --- | --- |
|  | OR | p | OR | P | OR | p | OR | P | OR | P |  |
| depression | **1.77** | **8.44e-23** | **0.95** | **7.64e-5** | **0.98** | **6.35e-6** | 1.01 | 0.0572 | 1.09 | 0.198 | 0.592 |
| irritability | **0.77** | **2.36e-6** | **0.94** | **1.06e-5** | **0.98** | **3.85e-9** | **1.02** | **0.000452** | 1.06 | 0.361 | 0.570 |
| psychosis | 0.96 | 0.684 | 1.02 | 0.313 | 1.01 | 0.334 | **1.06** | **2.77e-14** | 0.88 | 0.242 | 0.599 |
| VAB | **0.70** | **1.04e-9** | 1.00 | 0.863 | **0.99** | **7.95e-5** | **1.04** | **1.32e-12** | 0.97 | 0.630 | 0.601 |
| apathy | 1.04 | 0.482 | 1.02 | 0.156 | 1.00 | 0.900 | **1.04** | **9.84e-15** | 1.06 | 0.411 | 0.572 |
| POB | 0.99 | 0.865 | 1.02 | 0.0603 | 1.00 | 0.322 | **1.05** | **1.28e-19** | **1.52** | **5.72e-10** | 0.586 |
| Cognitive impairment | 1.10 | 0.0814 | **1.05** | **8.75e-5** | 1.01 | 0.112 | **1.10** | **2.08e-54** | 0.96 | 0.540 | 0.638 |

AUC represents the area under the curve, and is a measure of the ability of the covariates to jointly predict symptom presence. VAB = violent/aggressive behaviour. POB= perseverative/obsessive behaviour. Analysis applied to 5854 individuals with at least one recorded symptom endorsement (not necessarily positive). All variables fitted simultaneously. Significant associations shown in **bold**

Table 4 lists the PRS-symptom associations reaching nominal (p<0.05) significance, with only the PRS cutoff giving the most significant result being shown. Full PRS-symptom analyses are shown in **Supplementary Tables 2-10.** The schizophrenia PRS showed significant associations with psychosis, irritability and violent and aggressive behaviour after correction for both the number of PRS-symptom tests and the six PRS cutoffs, as did the MDD PRS with depression, irritability and violent and aggressive behaviour. The associations of schizophrenia PRS with depression and perseverative/obsessive behaviour were significant after correction for the number of PRS-symptom tests, as was the association between BPD PRS and depression. Of the primary hypotheses, the following nominally significant associations were also observed: MDD PRS with apathy, ADHD PRS with violent and aggressive behaviour, PD PRS with cognitive impairment and (interestingly, given the relatively small training GWAS) OCD PRS with perseverative/obsessive behaviour. In each case, increased PRS was associated with an increased risk of the symptom (as shown by odds ratios >1 in **Table 4**). Associations of the intelligence PRS with violent and aggressive behaviour, apathy and cognitive impairment were significant after correction for both the number of PRS-symptom tests and the 6 PRS cutoffs, while the association between intelligence PRS and irritability was significant after correction for the number of PRS-symptom tests. The associations with intelligence PRS have OR <1 in Table 2, indicating that decreased PRS (*i.e*. reduced intelligence) is associated with increased risk of the symptom. In general, PRS-symptom associations in **Table 4** surviving correction for multiple testing of symptom-PRS comparisons showed at least nominally significant association over a range of PRS cutoffs **(Figures 1, 2 and 3, Supplementary Tables 2**,**3**,**10**) thus increasing confidence that these results are robust. Conversely, the association between PD PRS and cognitive impairment was only significant for PRS cutoff p<0.0001 (**Supplementary Table 9**), making it likely that this is a false positive. This is also the case for the apparent association between psychosis and AD PRS (**Supplementary Table 8**). It can be seen from **Table 4** that the proportion of symptom variance accounted for by the PRS, as measured by the Nagelkerke R^2^, is small (<1%). Likewise, the ability of the PRS to predict symptom presence, as measured by the AUC, is limited (AUCs <0.55). For comparison, the Psychiatric Genomics Consortium (9) observed that schizophrenia PRS accounted for ∼7% of variance in schizophrenia liability, with AUC around 0.7. AUC of 0.8 is generally required for a clinically useful predictor (30).

Since sex, CAG length, age at motor onset and disease duration were found to correlate significantly with symptom risk (**Table 3**), the association analyses between PRS and symptom were repeated conditioning on the effects of the factors found to be significantly associated with symptom presence. This made little difference to the significance of the PRS-symptom associations (results not shown). Together, these factors give AUC of 0.57-0.64 (**Table 3**), and adding the PRS makes little difference to this (results not shown).

**Table 4.** HD symptom-polygenic risk score associations reaching nominal significance.

| SYMPTOM | Risk score | PRS CUT-OFF | OR | R<sup>2</sup> (%) | AUC | P VALUE |
| --- | --- | --- | --- | --- | --- | --- |
| Depression | **MDD** | **<u>0.01</u>** | **<u>1.17</u>** | **<u>0.78</u>** | **<u>0.547</u>** | **<u>9.01 x 10<sup>-8</sup></u>** |
|  | *Schizophrenia* | *0.05* | *1.12* | *0.37* | *0.528* | *2.47 x 10<sup>-4</sup>* |
|  | *BPD* | *0.01* | *1.11* | *0.36* | *0.532* | *3.06 x 10<sup>-4</sup>* |
|  | Intelligence | 0.001 | 0.93 | 0.18 | 0.519 | 9.51 x 10<sup>-3</sup> |
| Irritability | **Schizophrenia** | **<u>0.05</u>** | **<u>1.17</u>** | **<u>0.76</u>** | **<u>0.542</u>** | **<u>1.03 x 10<sup>-7</sup></u>** |
|  | **MDD** | **<u>0.001</u>** | **<u>1.12</u>** | **<u>0.45</u>** | **<u>0.535</u>** | **<u>4.42 x 10<sup>-5</sup></u>** |
|  | *Intelligence* | *1* | *0.90* | *0.37* | *0.531* | *1.99 x 10<sup>-4</sup>* |
| Psychosis | **Schizophrenia** | **<u>0.001</u>** | **<u>1.20</u>** | **<u>0.64</u>** | **<u>0.548</u>** | **<u>5.98 x 10<sup>-5</sup></u>** |
|  | MDD | 0.05 | 1.14 | 0.34 | 0.537 | 3.40 x 10<sup>-3</sup> |
|  | Alzheimer's | 0.001 | 1.10 | 0.19 | 0.529 | 0.0265 |
|  | Intelligence | 0.0001 | 0.90 | 0.24 | 0.534 | 0.0147 |
| VAB | **Intelligence** | **<u>0.01</u>** | **<u>0.89</u>** | **<u>0.50</u>** | **<u>0.537</u>** | **<u>2.44 x 10<sup>-5</sup></u>** |
|  | **Schizophrenia** | **<u>0.05</u>** | **<u>1.13</u>** | **<u>0.46</u>** | **<u>0.532</u>** | **<u>5.46 x 10<sup>-5</sup></u>** |
|  | **MDD** | **<u>0.001</u>** | **<u>1.13</u>** | **<u>0.44</u>** | **<u>0.536</u>** | **<u>8.45 x 10<sup>-5</sup></u>** |
|  | *BPD* | *0.001* | *1.11* | *0.34* | *0.527* | *5.35 x 10<sup>-4</sup>* |
|  | <u>ADHD</u> | <u>0.05</u> | <u>1.09</u> | <u>0.22</u> | <u>0.523</u> | <u>5.26 x 10<sup>-3</sup></u> |
|  | ASD | 0.01 | 1.06 | 0.11 | 0.517 | 0.0451 |
| POB | *Schizophrenia* | *0.01* | *1.12* | *0.39* | *0.531* | *1.47 x 10<sup>-4</sup>* |
|  | BPD | 0.5 | 1.08 | 0.18 | 0.521 | 9.25 x 10<sup>-3</sup> |
|  | Intelligence | 0.01 | 0.93 | 0.18 | 0.521 | 0.0103 |
|  | MDD | 0.5 | 1.07 | 0.15 | 0.519 | 0.0188 |
|  | <u>OCD</u> | <u>0.001</u> | <u>1.07</u> | <u>0.15</u> | <u>0.521</u> | <u>0.0190</u> |
| Apathy | **Intelligence** | **<u>1</u>** | **<u>0.90</u>** | **<u>0.41</u>** | **<u>0.534</u>** | **<u>9.27 x 10<sup>-5</sup></u>** |
|  | <u>MDD</u> | <u>0.05</u> | <u>1.08</u> | <u>0.18</u> | <u>0.521</u> | <u>9.17 x 10<sup>-3</sup></u> |
|  | BPD | 0.001 | 1.07 | 0.14 | 0.519 | 0.0222 |
|  | <u>Schizophrenia</u> | <u>0.0001</u> | <u>1.07</u> | <u>0.13</u> | <u>0.518</u> | <u>0.0254</u> |
|  | ASD | 0.001 | 1.06 | 0.10 | 0.516 | 0.0499 |
| Cognitive impairment | **Intelligence** | **<u>0.01</u>** | **<u>0.89</u>** | **<u>0.45</u>** | **<u>0.535</u>** | **<u>3.90 x 10<sup>-5</sup></u>** |
|  | <u>Parkinson's</u> | <u>0.0001</u> | <u>1.09</u> | <u>0.26</u> | <u>0.524</u> | <u>1.67 x 10<sup>-3</sup></u> |
|  | MDD | 0.5 | 1.09 | 0.23 | 0.524 | 3.51 x 10<sup>-3</sup> |

VAB= violent/aggressive behaviour, POB=perserverative/obsessive behaviour. MDD= major depressive disorder. BPD= bipolar disorder. ASD= autism spectrum disorder. ADHD = attention deficit hyperactivity disorder. R<sup>2</sup>= Nagelkerke R<sup>2</sup>. OR= Odds ratio for presence of symptom associated with 1 s.d. increase in PRS.
Results in *italics* satisfy the Bonferroni corrected p-value cut-off 7.94 x 10<sup>-4</sup> (corrected for 63 possible PRS-phenotype comparisons), and results in **bold** satisfy the Bonferroni corrected p-value cut of 1.32 x 10<sup>-4</sup> (corrected for the 63 PRS-symptom comparisons and 6 PRS cut-offs). Primary symptom-PRS hypotheses (see text) are underlined.

**Figure 2.**
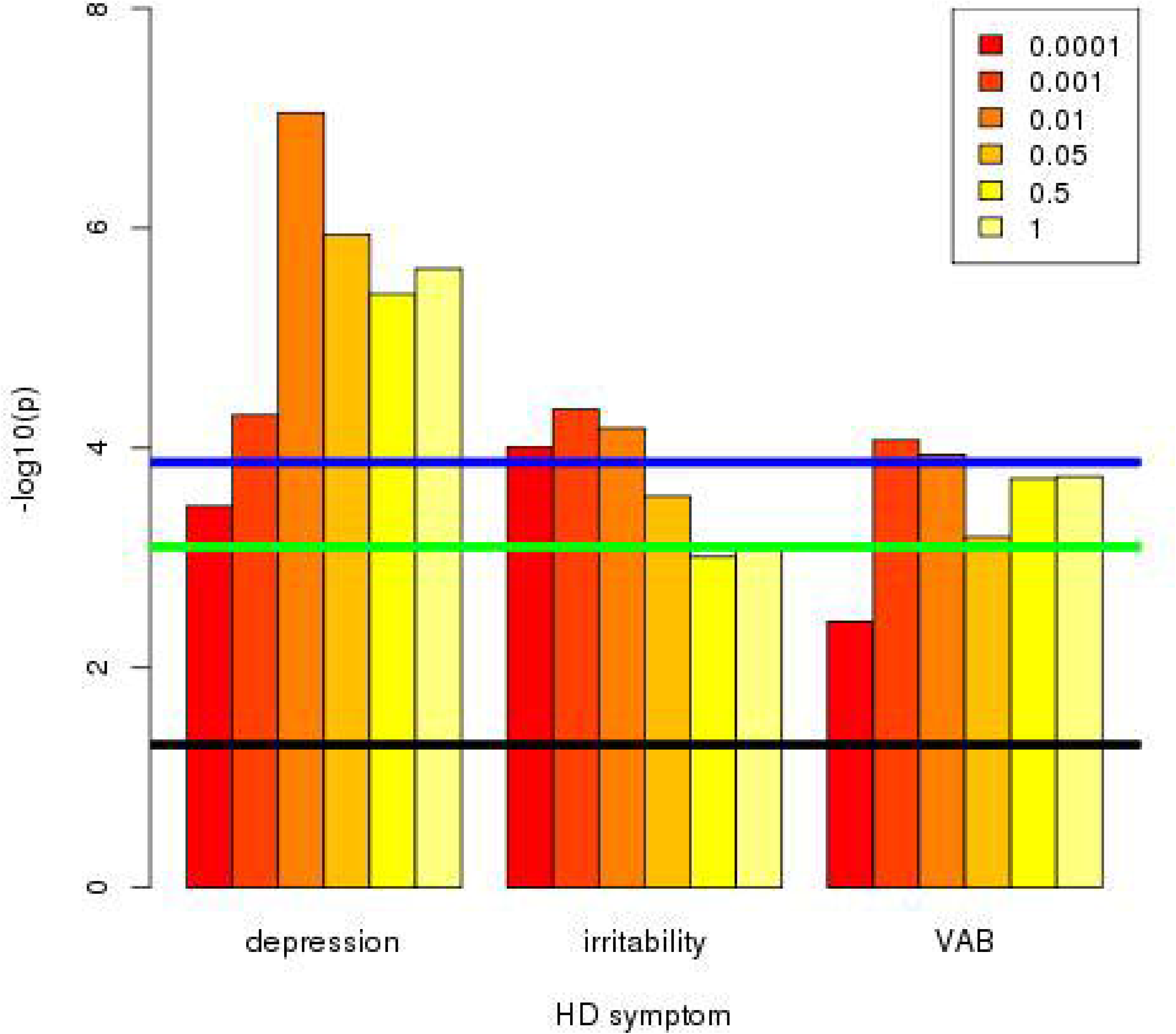
Association of increased major depression PRS with increased HD symptom frequency. Numbers in the box correspond to the p-value thresholds used to derive the PRS. Black line corresponds to nominally significant association (p=0.05). Green line indicates associations significant after Bonferroni correction for 63 PRS-symptom comparisons (9 PRS x 7 symptoms). Blue line indicates associations significant after Bonferroni correction for 63 PRS-symptom comparisons and 6 PRS cutoffs. Note: Only symptoms with at least one p-value reaching the green line are shown.

**Figure 3.**
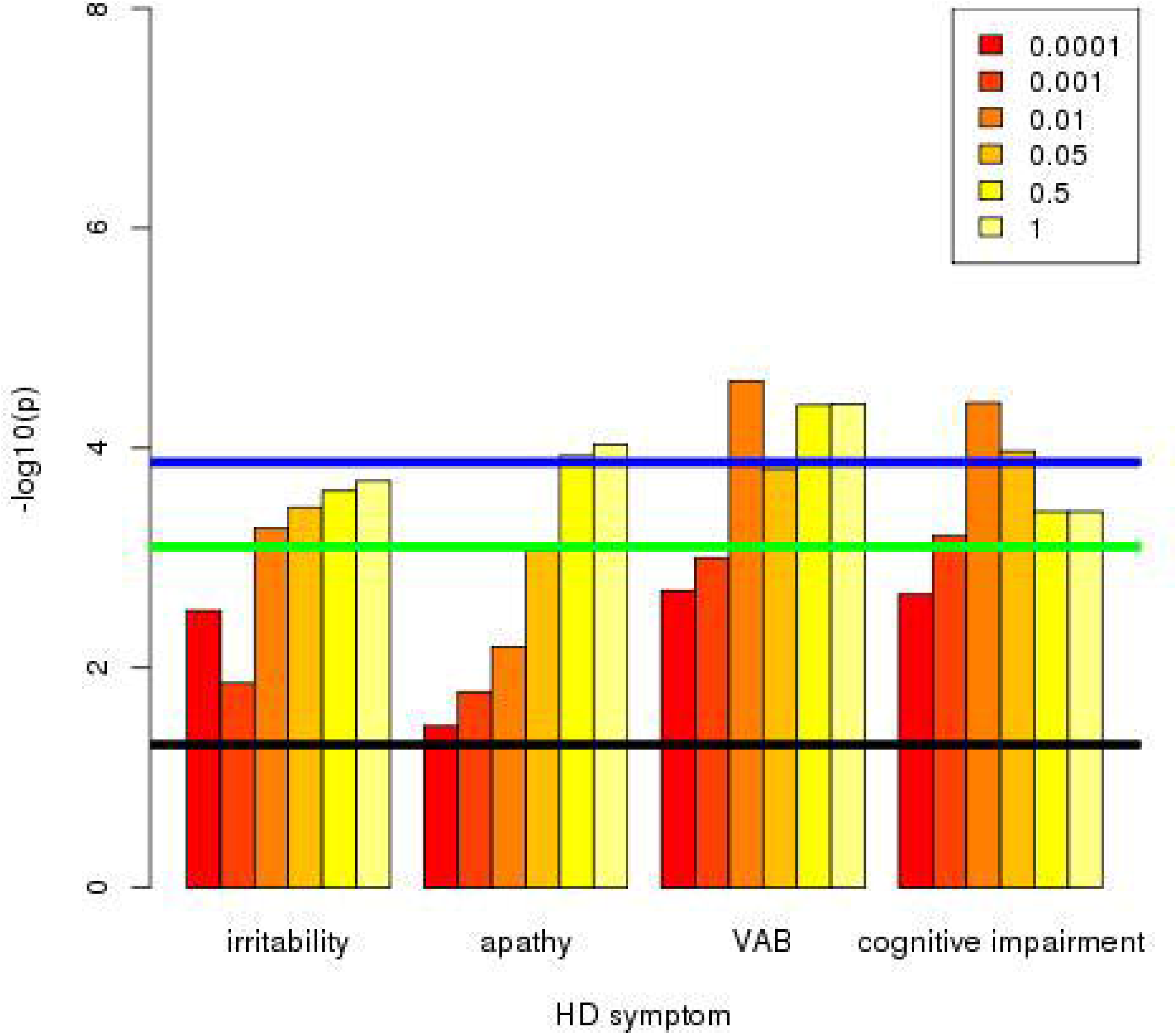
Association of decreased intelligence PRS with increased HD symptom frequency. Numbers in the box correspond to the p-value thresholds used to derive the PRS. Black line corresponds to nominally significant association (p=0.05). Green line indicates associations significant after Bonferroni correction for 63 PRS-symptom comparisons (9 PRS x 7 symptoms). Blue line indicates associations significant after Bonferroni correction for 63 PRS-symptom comparisons and 6 PRS cutoffs. Note: Only symptoms with at least one p-value reaching the green line are shown.

For symptoms with a significant association with sex (depression, irritability and violent and aggressive behaviour) and PRS with a significant association in the whole sample, the PRS-symptom association analyses were run in males and females separately (**Supplementary Table 11**). There were no significant differences in OR between males and females.

PRS for schizophrenia, MDD, BPD and intelligence are associated with multiple symptoms in **Table 4**. Since these symptoms are correlated (**Supplementary Table 1**), it is unclear which symptom is driving the association. Likewise, symptoms are often associated with multiple PRS in **Table 4**. These PRS will be correlated, due to genetic overlap between psychiatric disorders (13). We therefore performed logistic regression of each symptom on all the significant PRS from **Table 4** simultaneously, including the six other symptom diagnoses as covariates (**Supplementary Table 12**). This analysis highlighted specific associations between MDD PRS and depression, schizophrenia PRS and both psychosis and irritability, BPD PRS and violent/aggressive behaviour and intelligence PRS and cognitive decline. AD PRS and PD PRS were significantly associated with psychosis and cognitive impairment respectively, but these are likely to be false positives, as discussed above.

The number of symptoms seen in any individual can be regarded as a surrogate for disease severity, and might be expected to correlate with PRS. A linear regression of PRS on symptom count (0-7) was therefore performed, (**Supplementary Table 13**). Significant associations with symptom count were observed for schizophrenia (p=4.45×10^-9^), intelligence (p=5.78×10^-8^), MDD (p=4.17×10^-8^) and BD (p=0.00268), with the latter being attributable to correlation with the other PRS. To test whether association of PRS with individual symptoms can be explained by the general association with symptom count, the number of other symptoms was included as a covariate in the regression of PRS on symptom presence. Psychosis and irritability were found to be associated with schizophrenia PRS independently of other symptoms (**Supplementary Table 14**). Depression was associated with MDD PRS across all cutoffs (**Supplementary Table 15**), as was cognitive impairment with intelligence PRS (**Supplementary Table 16**). These results corroborate the specific PRS-symptom associations from the previous paragraph.

## Discussion

We show significant genetic overlaps between psychiatric disorders and psychiatric symptoms in HD, along with genetic overlap between intelligence and cognitive symptoms in HD. There is little overlap between the two neurodegenerative disorders and neuropsychiatric symptoms or cognition in HD. While the overall pattern of individual associations is complex, reflecting the genetic relationships between the disorders, (see **Figure 4** for a graphical overview), we were able to identify that MDD PRS is specifically associated with depression, schizophrenia PRS with psychosis and irritability, BPD PRS with violent/aggressive behaviour and intelligence PRS with cognitive impairment.

**Figure 4.**
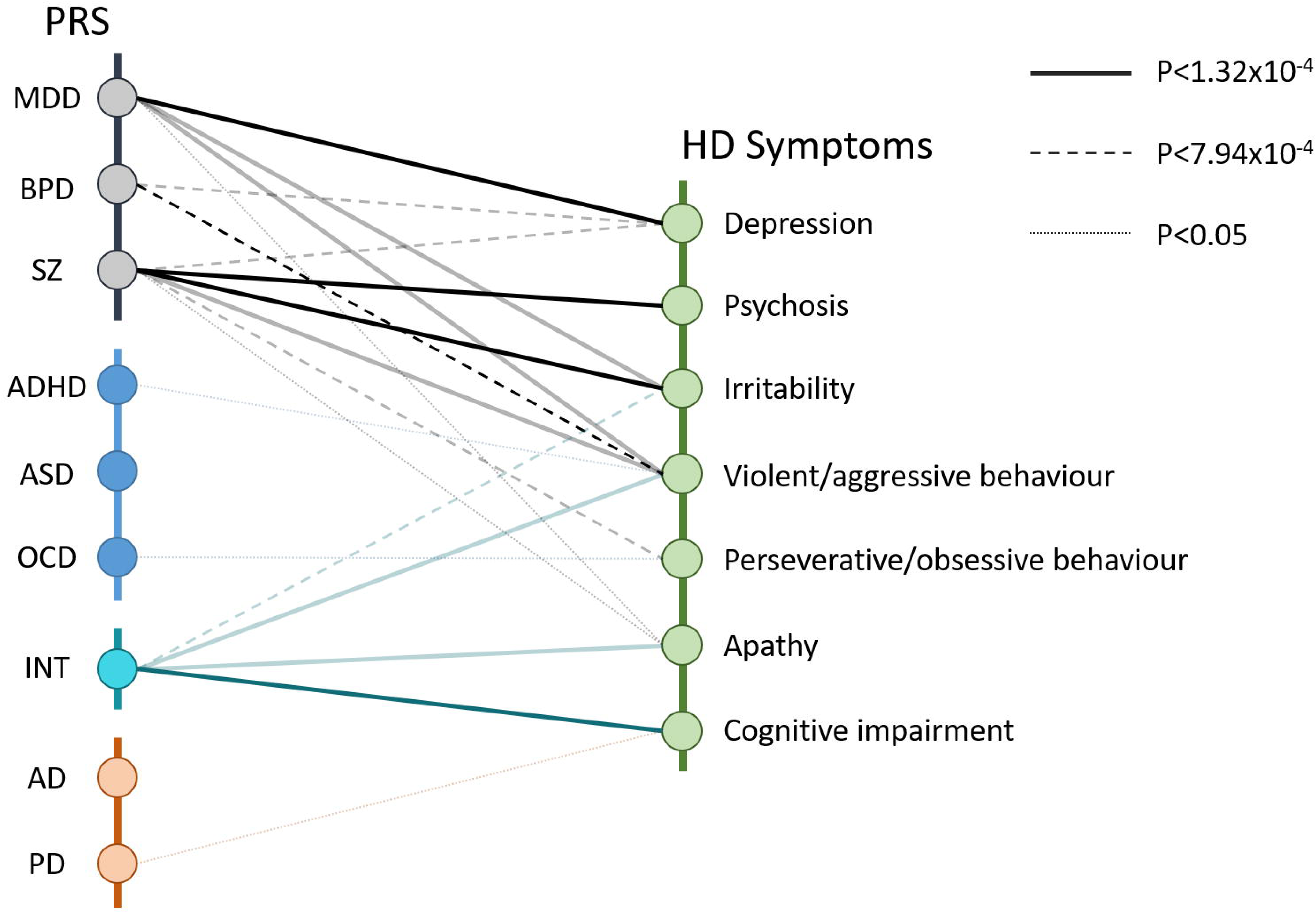
Pattern of association between HD symptoms and PRS from psychiatric, neurodegenerative and cognitive disorders. PRS grouped into psychiatric (black), neurodevelopmental (blue), neurodegenerative (orange) and cognitive (cyan) disorders. Solid lines show associations significant after correcting for 63 PRS-symptom combinations (9 PRS x 7 symptoms) and 6 PRS cutoffs (p<1.32×10^-4^). Dashed lines show associations significant after correcting for 63 PRS-symptom combinations (p<7.94×10^-4^). Dotted lines show nominally significant associations (p<0.05) in PRS-symptom combinations that were part of the primary analysis. Bold lines show PRS – symptom associations that are significant after correcting for other PRS and symptoms (Supplementary Table 12)

Our results are consistent with observations of shared genetic risk amongst psychiatric disorders but no overlap between the individual neurological disorders and no overlap with neuropsychiatric symptom risk in neurodegenerative disorders (13, 21, 29). By contrast, common genetic variation is associated with behavioural phenotypes (such as autistic behaviour and developmental delay) in severe monogenic neurodevelopmental disorders, with similar effect sizes (R^2^ of 0.6-0.8%) to those observed here (31). These results, along with ours, indicate that even in diseases previously assumed to be entirely attributable to single genetic variants, there is a contribution of polygenic risk in influencing phenotypic presentation (31).

Schizophrenia PRS is associated with psychosis (and irritability) independently of other symptoms and is the only PRS to predict HD psychosis. While psychosis in HD is around ten times more common than in the general population, it is seen in only a minority of cases (2), as in the participants studied here (11%). There are also reports of clustering of psychotic symptoms in HD families (6, 7), implying a genetic contribution to these symptoms. Previously, Tsuang *et al.* (8) tested for association of psychosis in HD with a set of 214 SNPs chosen from candidate genes for schizophrenia, HD or psychosis in neurodegenerative disorders. None of their associations survived correction for multiple testing, possibly due to the small sample size (47 HD cases with psychosis, 126 without psychosis). Of the SNPs tested by Tsuang *et al.*, 183 were available in our data; a polygenic risk score generated from these (with effect sizes taken from the PGC schizophrenia GWAS) showed no association with psychosis in our HD sample (OR=0.97, p=0.536). The highly significant association between schizophrenia PRS and psychosis observed in our sample shows the benefits of a large sample size of HD patients, along with a larger set of SNPs systematically selected for association with schizophrenia in a powerful schizophrenia GWAS. The schizophrenia PRS is also the most significantly associated with perseveration, one of the most characteristic behavioural problems in HD. The PRS for OCD also influences the presence of perseverative/obsessive behaviour **(Table 4, Supplementary Table 12)**, despite the OCD PRS having been derived from a much smaller and less powerful sample than that for schizophrenia.

Schizophrenia may be regarded as a neurodevelopmental disorder with origins in foetal development that manifest in symptoms most usually in early adulthood (32), whereas psychotic symptoms in HD generally manifest later in life (2, 33). The genes that contribute to the schizophrenia PRS are preferentially expressed in the medium spiny neurons of the striatum (34), which are the most vulnerable cell types in HD: by the time HD motor symptoms are manifest up to half of these neurons have died (35). Susceptibility to psychiatric symptoms in surviving cells is influenced by the schizophrenia polygenic risk, and provides a potential explanation for the increased rates of psychotic symptoms observed in HD patients.

Cognitive decline and dementia are part of HD progression, with executive and psychomotor function often the first deficits noted, with memory problems later in disease (2). Longer CAG repeat tracts are associated with increased risk of cognitive deficits in our sample: the previous data on this relationship have been inconsistent (2, 36). Recent studies have shown that poorer performance in the symbol digit modality test was associated with longer *HTT* CAG repeat lengths (37), and poorer cognitive function was seen with increasing repeat lengths in the disease causing range (38). The clinical characteristics questionnaire used to assess symptoms for this study seeks an integrated measure of cognitive decline by asking about problems that might interfere with performing everyday functions and is relatively crude, though this will be partly offset by the larger size of our study (1000s rather than 100s of participants).

It is notable that neither apathy nor cognitive dysfunction in HD are correlated with PRS for psychiatric or neurodegenerative disorders apart from nominal associations between apathy and MDD, schizophrenia and BPD PRS, which become non-significant when accounting for the presence of other symptoms (see **Supplementary Table 12**). However, both are significantly correlated with a PRS measuring intelligence. This suggests that apathy in HD should be considered as a cognitive, rather than psychiatric, symptom (unlike depression), and is also consistent with apathy being the only psychiatric symptom to correlate with disease progression (39). Higher PRS for intelligence is associated with later cognitive decline in HD, as it is in AD (14). The genes associated with intelligence are highly expressed in medium spiny neurons of the striatum and pyramidal neurons of the CA1 hippocampus (14), early targets of degeneration in HD and AD respectively, potentially contributing to the severe cognitive decline seen in these diseases. Surprisingly, there is no significant association between the presence of cognitive deficits in HD and genetic risk for AD, though AD PRS predicts memory decline and poorer cognitive performance in healthy children and adults well before the age of risk for AD (40, 41). In children, *HTT* CAG repeat length itself shows a J-shaped relationship with cognition, with maximum cognition at 40-41 repeats (38). Since higher intelligence is associated with better health and increased well-being (42), there may thus be a selective advantage of longer *HTT* CAG repeat length, below the threshold for HD, in the wider population.

Depression is very common in HD and more common in females (2). MDD PRS is specifically associated with increased risk of depression in HD, and the converse is also true (**Supplementary Table 12**). However, there were no significant sex differences in these associations (**Supplementary Table 11**). There is a significant correlation in our sample of increased likelihood of depression with shorter CAG repeat lengths in the disease-causing allele in *HTT* (**Table 3**), along with an association with earlier age at motor onset. The reason for these associations is unclear; the obvious explanation that longer disease duration makes depression more likely is not the case in our sample (**Table 3**). In fact, the apparent associations between depression and both age at motor onset and CAG repeat length are explained by the residual of age at motor onset after correction for CAG length (see (5) for how this is derived). Participants with an earlier than expected age at onset (given their CAG length) are more likely to have depression. This could be due to increased environmental stress among individuals with earlier motor onset, and also that some genetic modifiers of age at motor onset may also be associated with depression in HD. The few previous studies, each examining less than 100 participants, detected no relationship between CAG length and presence of depression (43–45). In the non-disease causing range, longer HTT CAG length from 24 to 38 repeats was associated with an increased likelihood of depression (46).

Irritability, like depression, is very common in HD and has a significantly reduced risk with longer CAG length (**Table 3**) but unlike depression, longer duration of disease makes irritability more likely. This association is not explained by increased age. Schizophrenia PRS is associated with increased risk of irritability (**Table 4, Supplementary Table 12**). Violent/aggressive behaviour can be considered to be an extreme manifestation of irritability, and is associated with PRS for bipolar disorder (**Supplementary Table 12**), with the apparently more significant associations with schizophrenia, intelligence and MDD PRS (**Table 4**) attributable to correlations between violent/aggressive behaviour and other symptoms. As in the wider population violence and aggressive behaviour is more common in men (2).

The study presented here has a number of limitations. As noted, the clinical characteristics questionnaire is a relatively crude instrument. The larger size of this study compared with previous studies partly mitigates its limitations, and these initial findings provide a platform for more detailed studies using specific psychiatric instruments. We note that all Enroll-HD participants now undergo a short problem behaviours assessment battery (PBA), which may account for the increased frequency of perseverative/obsessive behaviour observed in Enroll-HD relative to REGISTRY. It would be useful to investigate further the predictive power of the PRS using more detailed clinical data, including age at onset, on the psychiatric symptoms in HD. It would also be interesting to explore the overlap of multiple symptoms and the psychiatric and cognitive PRS to attempt to establish directions of causation.

It is notable that the PRS for psychiatric diseases associate with increased risk of developing parallel phenotypic behavioural and psychiatric symptoms in HD participants. The data available in the ongoing Enroll-HD study will provide for a much more detailed analyses of these symptoms, their aetiology and treatment and may in turn inform the psychiatric diseases. The lack of genetic overlap with other neurodegenerative disorders, consistent with the Brainstorm study (13), implies different underlying pathways leading to degeneration in the different neurodegenerations that may relate to the disease-specific characteristic differential neuronal vulnerabilities to degeneration. Striatial medium spiny neurons are most vulnerable in HD, dopaminergic neurons of the substantia nigra in PD and hippocampal and entorhinal cortical neurons in AD. The differential vulnerability may relate to pathways essential to the survival and continued function of each specific cell type and these are likely to be different in different neurodegenerations (47). Thus, psychiatric symptoms may be mediated by common dysfunctional pathways in surviving cells whereas cognitive and neurodegenerative symptoms are likely due to specific regional cellular populations degenerating via different pathways. These data provide a rationale for treatments used in the wider population for depression and psychotic symptoms, to be used for these symptoms in HD.

## Acknowledgements

This work was supported by the CHDI Foundation, the National Institutes of Health USA (U01NS082079, R01NS040068, R01NS091161, P50NS016367, and R01NS049206) and the Medical Research Council (UK; MR/L010305/1). Dr. Massey received a fellowship from the MRC (MR/P001629/1) and Mr. McAllister received a studentship from the Cardiff University School of Medicine. The Enroll-HD and REGISTRY, studies would not be possible without the vital contribution of the research participants and their families. Individuals who contributed to the collection subject data can be found at https://www.enroll-hd.org/acknowledgments/ for Enroll-HD, and in Supplementary Information Appendix B for REGISTRY. Members of the EHDN Behavioral Phenotype Working Group are listed in Supplementary Information Appendix C.

