## Supplementary Information for "Genetic risk underlying psychiatric and cognitive symptoms in Huntington’s Disease"

**Supplementary Figure 1:**

Flow diagram showing how the sample used to test association between PRS and symptoms was derived.

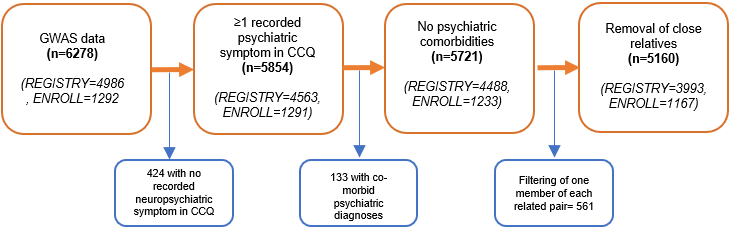

**Supplementary Table 1**

Correlations between symptoms.

|  | irb | psy | VAB | cog | apt | POB |
| --- | --- | --- | --- | --- | --- | --- |
| dep | 0.242 | 0.139 | 0.184 | 0.168 | 0.291 | 0.161 |
| irb |  | 0.162 | 0.466 | 0.194 | 0.270 | 0.303 |
| psy |  |  | 0.250 | 0.165 | 0.139 | 0.199 |
| VAB |  |  |  | 0.213 | 0.234 | 0.281 |
| cog |  |  |  |  | 0.300 | 0.210 |
| apt |  |  |  |  |  | 0.278 |

All correlations highly significant (p<2.2x10^-16^)

dep = depression, irb = irritability, psy = psychosis, VAB = violent/aggressive behaviour. POB= perseverative/obsessive behaviour. cog=cognitive impairment, apt=apathy

**Supplementary Table 2:** All PRS-symptom analyses for SCZ PRS (no MHC)

| Symptom | Statistic | PRS cutoff | | | | | |
| --- | --- | --- | --- | --- | --- | --- | --- |
|  |  | 0.0001 | 0.001 | 0.01 | 0.05 | 0.5 | 1 |
| dep | OR | 1.07(1.01,1.14) | 1.08(1.02,1.15) | 1.10(1.04,1.17) | 1.12(1.05,1.18) | 1.10(1.04,1.17) | 1.10(1.04,1.17) |
|  | R2 | 0.0015 | 0.0020 | 0.0030 | 0.0037 | 0.0029 | 0.0029 |
|  | p | 0.0193 | 0.00715 | 0.000929 | 0.000247 | 0.00114 | 0.00115 |
| irb | OR | 1.09(1.03,1.16) | 1.11(1.05,1.17) | 1.16(1.09,1.22) | 1.17(1.10,1.24) | 1.15(1.09,1.22) | 1.15(1.08,1.21) |
|  | R2 | 0.0025 | 0.0035 | 0.0067 | 0.0076 | 0.0061 | 0.0060 |
|  | p | 0.00213 | 0.000318 | 5.60e-7 | 1.03e-7 | 1.82e-6 | 2.31e-6 |
| psy | OR | 1.18(1.09,1.29) | 1.20(1.10,1.31) | 1.19(1.09,1.30) | 1.17(1.08,1.28) | 1.18(1.08,1.29) | 1.17(1.08,1.28) |
|  | R2 | 0.0057 | 0.0064 | 0.0060 | 0.0051 | 0.0053 | 0.0051 |
|  | p | 0.000148 | 5.98e-5 | 0.000103 | 0.000342 | 0.000260 | 0.000350 |
| apt | OR | 1.07(1.01,1.13) | 1.05(0.99,1.11) | 1.06(1.00,1.12) | 1.05(0.99,1.11) | 1.03(0.98,1.09) | 1.03(0.98,1.09) |
|  | R2 | 0.0013 | 0.0008 | 0.0010 | 0.0007 | 0.0004 | 0.0004 |
|  | p | 0.0254 | 0.0858 | 0.0549 | 0.0936 | 0.231 | 0.245 |
| VAB | OR | 1.05(0.99,1.12) | 1.05(0.99,1.12) | 1.11(1.05,1.18) | 1.13(1.07,1.20) | 1.11(1.04,1.18) | 1.11(1.04,1.18) |
|  | R2 | 0.0008 | 0.0008 | 0.0035 | 0.0046 | 0.0032 | 0.0032 |
|  | p | 0.0927 | 0.0892 | 0.000432 | 5.46e-5 | 0.000807 | 0.000750 |
| POB | OR | 1.08(1.02,1.14) | 1.11(1.05,1.18) | 1.12(1.06,1.18) | 1.10(1.04,1.16) | 1.12(1.05,1.18) | 1.12(1.05,1.18) |
|  | R2 | 0.0019 | 0.0037 | 0.0039 | 0.0027 | 0.0037 | 0.0038 |
|  | p | 0.00799 | 0.000246 | 0.000147 | 0.00161 | 0.000215 | 0.000208 |
| cog | OR | 1.01(0.95,1.07) | 1.03(0.97,1.08) | 1.05(1.00,1.11) | 1.06(1.00,1.12) | 1.04(0.98,1.10) | 1.04(0.98,1.10) |
|  | R2 | 2.87e-5 | 0.0002 | 0.0009 | 0.0009 | 0.0005 | 0.0005 |
|  | p | 0.742 | 0.384 | 0.0678 | 0.0592 | 0.164 | 0.152 |

dep = depression, irb = irritability, psy = psychosis, apt = apathy, VAB = violent/aggressive behaviour. POB= perseverative/obsessive behaviour. cog=cognitive impairment.

**Supplementary Table 3:** All PRS-symptom analyses for MDD PRS

| Symptom | Statistic | PRS cutoff | | | | | |
| --- | --- | --- | --- | --- | --- | --- | --- |
|  |  | 0.0001 | 0.001 | 0.01 | 0.05 | 0.5 | 1 |
| dep | OR | 1.11(1.05,1.18) | 1.13(1.06,1.19) | 1.17(1.11,1.24) | 1.16(1.09,1.22) | 1.15(1.08,1.22) | 1.15(1.09,1.22) |
|  | R2 | 0.0035 | 0.0045 | 0.0078 | 0.0064 | 0.0058 | 0.0061 |
|  | p | 0.000339 | 5.04e-5 | 9.01e-8 | 1.14e-6 | 3.97e-6 | 2.34e-6 |
| irb | OR | 1.12(1.06,1.18) | 1.12(1.06,1.19) | 1.12(1.06,1.19) | 1.11(1.05,1.17) | 1.10(1.04,1.16) | 1.10(1.04,1.16) |
|  | R2 | 0.0041 | 0.0045 | 0.0043 | 0.0036 | 0.0029 | 0.0030 |
|  | p | 9.85e-5 | 4.42e-5 | 6.67e-5 | 0.000274 | 0.000985 | 0.000823 |
| psy | OR | 1.08(0.99,1.18) | 1.13(1.03,1.23) | 1.13(1.03,1.23) | 1.14(1.04,1.24) | 1.12(1.03,1.22) | 1.11(1.02,1.21) |
|  | R2 | 0.0012 | 0.0028 | 0.0029 | 0.0034 | 0.0026 | 0.0022 |
|  | p | 0.0785 | 0.00744 | 0.00655 | 0.00340 | 0.0109 | 0.0162 |
| apt | OR | 1.02(0.96,1.07) | 1.04(0.98,1.10) | 1.06(1.00,1.12) | 1.08(1.02,1.14) | 1.07(1.01,1.13) | 1.07(1.01,1.13) |
|  | R2 | 7.81e-5 | 0.0005 | 0.0012 | 0.0018 | 0.0016 | 0.0016 |
|  | p | 0.588 | 0.160 | 0.0356 | 0.00917 | 0.0157 | 0.0149 |
| VAB | OR | 1.09(1.03,1.16) | 1.13(1.06,1.20) | 1.13(1.06,1.20) | 1.11(1.05,1.18) | 1.12(1.06,1.19) | 1.12(1.06,1.19) |
|  | R2 | 0.0023 | 0.0044 | 0.0042 | 0.0033 | 0.0039 | 0.0040 |
|  | p | 0.00379 | 8.45e-5 | 0.000116 | 0.000652 | 0.000189 | 0.000183 |
| POB | OR | 1.00(0.95,1.06) | 1.02(0.97,1.08) | 1.05(0.99,1.11) | 1.06(1.00,1.12) | 1.07(1.01,1.13) | 1.07(1.01,1.13) |
|  | R2 | 7.51e-6 | 0.0002 | 0.0007 | 0.0010 | 0.0015 | 0.0013 |
|  | p | 0.868 | 0.400 | 0.106 | 0.0570 | 0.0188 | 0.0275 |
| cog | OR | 1.02(0.96,1.07) | 1.07(1.01,1.13) | 1.05(0.99,1.11) | 1.09(1.03,1.15) | 1.09(1.03,1.15) | 1.08(1.02,1.14) |
|  | R2 | 8.72e-5 | 0.0015 | 0.0008 | 0.0023 | 0.0023 | 0.0020 |
|  | p | 0.566 | 0.0167 | 0.0777 | 0.00354 | 0.00351 | 0.00546 |

dep = depression, irb = irritability, psy = psychosis, apt = apathy, VAB = violent/aggressive behaviour. POB= perseverative/obsessive behaviour. cog=cognitive impairment.

**Supplementary Table 4:** All PRS-symptom analyses for BPD PRS

| Symptom | Statistic | PRS cutoff | | | | | |
| --- | --- | --- | --- | --- | --- | --- | --- |
|  |  | 0.0001 | 0.001 | 0.01 | 0.05 | 0.5 | 1 |
| dep | OR | 1.04(0.98,1.10) | 1.07(1.00,1.13) | 1.11(1.05,1.18) | 1.11(1.05,1.18) | 1.07(1.01,1.14) | 1.07(1.01,1.14) |
|  | R2 | 0.0005 | 0.0012 | 0.0036 | 0.0034 | 0.0016 | 0.0016 |
|  | p | 0.198 | 0.0336 | 0.000306 | 0.000466 | 0.0161 | 0.0170 |
| irb | OR | 1.04(0.99,1.10) | 1.04(0.98,1.10) | 1.04(0.98,1.10) | 1.01(0.96,1.07) | 1.01(0.96,1.07) | 1.01(0.96,1.07) |
|  | R2 | 0.0006 | 0.0006 | 0.0005 | 4.20e-5 | 4.24e-5 | 4.11e-5 |
|  | p | 0.138 | 0.150 | 0.161 | 0.692 | 0.691 | 0.695 |
| psy | OR | 1.00(0.91,1.09) | 1.02(0.94,1.12) | 1.05(0.97,1.15) | 1.06(0.97,1.15) | 1.05(0.96,1.15) | 1.07(0.98,1.17) |
|  | R2 | 1.72e-6 | 0.0001 | 0.0006 | 0.0006 | 0.0005 | 0.0009 |
|  | p | 0.947 | 0.615 | 0.236 | 0.212 | 0.258 | 0.141 |
| apt | OR | 1.05(0.99,1.11) | 1.07(1.01,1.13) | 1.04(0.98,1.10) | 1.00(0.94,1.05) | 0.97(0.92,1.03) | 0.98(0.92,1.03) |
|  | R2 | 0.0006 | 0.0014 | 0.0005 | 3.44e-6 | 0.0002 | 0.0002 |
|  | p | 0.120 | 0.0222 | 0.185 | 0.909 | 0.358 | 0.436 |
| VAB | OR | 1.06(1.00,1.13) | 1.11(1.05,1.18) | 1.08(1.01,1.14) | 1.07(1.01,1.14) | 1.08(1.01,1.14) | 1.08(1.02,1.15) |
|  | R2 | 0.0012 | 0.0034 | 0.0017 | 0.0014 | 0.0016 | 0.0019 |
|  | p | 0.0413 | 0.000535 | 0.0152 | 0.0245 | 0.0175 | 0.00975 |
| POB | OR | 1.05(0.99,1.11) | 1.05(0.99,1.11) | 1.06(1.00,1.12) | 1.07(1.01,1.13) | 1.08(1.02,1.14) | 1.08(1.02,1.14) |
|  | R2 | 0.0008 | 0.0007 | 0.0010 | 0.0014 | 0.0018 | 0.0018 |
|  | p | 0.0839 | 0.118 | 0.0553 | 0.0254 | 0.00925 | 0.0113 |
| cog | OR | 0.98(0.92,1.03) | 1.00(0.95,1.06) | 1.02(0.97,1.08) | 1.03(0.97,1.09) | 1.03(0.98,1.09) | 1.03(0.97,1.09) |
|  | R2 | 0.0002 | 3.76e-7 | 0.0002 | 0.0003 | 0.0003 | 0.0003 |
|  | p | 0.421 | 0.970 | 0.451 | 0.305 | 0.275 | 0.323 |

dep = depression, irb = irritability, psy = psychosis, apt = apathy, VAB = violent/aggressive behaviour. POB= perseverative/obsessive behaviour. cog=cognitive impairment.

**Supplementary Table 5:** All PRS-symptom analyses for ADHD PRS

| Symptom | Statistic | PRS cutoff | | | | | |
| --- | --- | --- | --- | --- | --- | --- | --- |
|  |  | 0.0001 | 0.001 | 0.01 | 0.05 | 0.5 | 1 |
| dep | OR | 1.00(0.94,1.06) | 1.03(0.97,1.09) | 1.02(0.96,1.08) | 1.02(0.96,1.08) | 1.03(0.97,1.09) | 1.03(0.97,1.09) |
|  | R2 | 2.07e-7 | 0.0002 | 0.0001 | 0.0001 | 0.0002 | 0.0002 |
|  | p | 0.978 | 0.355 | 0.544 | 0.473 | 0.353 | 0.368 |
| irb | OR | 0.99(0.94,1.05) | 1.02(0.96,1.08) | 1.04(0.98,1.10) | 1.05(1.00,1.11) | 1.05(0.99,1.11) | 1.04(0.99,1.10) |
|  | R2 | 2.88e-5 | 0.0001 | 0.0005 | 0.0009 | 0.0007 | 0.0006 |
|  | p | 0.743 | 0.518 | 0.189 | 0.0741 | 0.100 | 0.135 |
| psy | OR | 1.00(0.91,1.09) | 1.04(0.95,1.13) | 0.98(0.90,1.07) | 1.05(0.96,1.14) | 1.07(0.98,1.16) | 1.06(0.97,1.16) |
|  | R2 | 1.94e-6 | 0.0003 | 9.51e-5 | 0.0004 | 0.0009 | 0.0007 |
|  | p | 0.944 | 0.369 | 0.624 | 0.302 | 0.135 | 0.173 |
| apt | OR | 0.99(0.94,1.05) | 1.01(0.96,1.07) | 1.03(0.97,1.09) | 1.03(0,98,1.09) | 1.02(0.97,1.08) | 1.02(0.97,1.08) |
|  | R2 | 2.70e-5 | 4.40e-5 | 0.0003 | 0.0004 | 0.0002 | 0.0001 |
|  | p | 0.750 | 0.684 | 0.324 | 0.239 | 0.432 | 0.477 |
| VAB | OR | 1.01(0.95,1.07) | 1.02(0.96,1.08) | 1.04(0.98,1.10) | 1.09(1.03,1.15) | 1.08(1.02,1.15) | 1.07(1.01.1.14) |
|  | R2 | 3.97e-5 | 8.54e-5 | 0.0004 | 0.0022 | 0.0018 | 0.0016 |
|  | p | 0.707 | 0.582 | 0.225 | 0.00526 | 0.0108 | 0.0179 |
| POB | OR | 1.01(0.95,1.07) | 0.98(0.93,1.04) | 1.01(0.96,1.07) | 1.00(0.95,1.06) | 0.99(0.94,1.05) | 0.98(0.93,1.04) |
|  | R2 | 3.06e-5 | 0.0001 | 4.73e-5 | 3.64e-6 | 3.31e-5 | 0.0001 |
|  | p | 0.738 | 0.525 | 0.677 | 0.908 | 0.728 | 0.478 |
| cog | OR | 0.97(0.92,1.03) | 0.99(0.93,1.04) | 1.03(0.98,1.09) | 1.02(0.97,1.08) | 1.05(1.00,1.11) | 1.05(1.00,1.11) |
|  | R2 | 0.0003 | 6.09e-5 | 0.0004 | 0.0002 | 0.0010 | 0.0009 |
|  | p | 0.324 | 0.632 | 0.237 | 0.414 | 0.0582 | 0.0691 |

dep = depression, irb = irritability, psy = psychosis, apt = apathy, VAB = violent/aggressive behaviour. POB= perseverative/obsessive behaviour. cog=cognitive impairment.

**Supplementary Table 6:** All PRS-symptom analyses for ASD PRS

| Symptom | Statistic | PRS cutoff | | | | | |
| --- | --- | --- | --- | --- | --- | --- | --- |
|  |  | 0.0001 | 0.001 | 0.01 | 0.05 | 0.5 | 1 |
| dep | OR | 0.98(0.92,1.04) | 0.98(0.93,1.04) | 0.99(0.93,1.05) | 0.99(0.93,1.05) | 0.97(0.92,1.03) | 0.97(0.92,1.03) |
|  | R2 | 0.0002 | 8.62e-5 | 6.29e-5 | 3.78e-5 | 0.0003 | 0.0003 |
|  | p | 0.446 | 0.574 | 0.631 | 0.710 | 0.304 | 0.291 |
| irb | OR | 1.00(0.94,1.05) | 1.03(0.97,1.08) | 1.01(0.95,1.07) | 1.00(0.95,1.06) | 1.00(0.95,1.06) | 0.99(0.94,1.05) |
|  | R2 | 6.72e-5 | 0.0002 | 1.72e-5 | 6.84e-8 | 8.63e-8 | 2.58e-5 |
|  | p | 0.874 | 0.383 | 0.800 | 0.987 | 0.986 | 0.756 |
| psy | OR | 1.02(0.93,1.11) | 1.04(0.95,1.13) | 1.06(0.97,1.16) | 1.04(0.95,1.13) | 1.05(0.96,1.14) | 1.06(0.97,1.16) |
|  | R2 | 7.18e-5 | 0.0003 | 0.0008 | 0.0002 | 0.0004 | 0.0007 |
|  | p | 0.670 | 0.396 | 0.168 | 0.434 | 0.300 | 0.184 |
| apt | OR | 1.03(0.97,1.09) | 1.06(1.00,1.12) | 1.03(0.97,1.08) | 1.01(0.96,1.07) | 1.00(0.94,1.05) | 0.99(0.94,1.05) |
|  | R2 | 0.0003 | 0.0010 | 0.0002 | 4.65e-5 | 8.93e-6 | 1.91e-5 |
|  | p | 0.328 | 0.0499 | 0.345 | 0.676 | 0.855 | 0.789 |
| VAB | OR | 1.01(0.95,1.07) | 1.06(1.00,1.12) | 1.06(1,00,1.13) | 1.03(0.98,1.10) | 1.03(0.97,1.09) | 1.03(0.97,1.09) |
|  | R2 | 4.77e-5 | 0.0010 | 0.0011 | 0.0004 | 0.0002 | 0.0002 |
|  | p | 0.681 | 0.0603 | 0.0451 | 0.255 | 0.379 | 0.366 |
| POB | OR | 0.96(0.91,1.02) | 1.04(0.98,1.10) | 1.02(0.97,1.08) | 1.01(0.96,1.07) | 1.01(0.96,1.07) | 1.01(0.95,1.07) |
|  | R2 | 0.0005 | 0.0004 | 0.0002 | 5.43e-5 | 5.67e-5 | 3.24e-5 |
|  | p | 0.159 | 0.219 | 0.397 | 0.656 | 0.648 | 0.730 |
| cog | OR | 0.99(0.94,1.05) | 1.05(1.00,1.11) | 1.03(0.98,1.09) | 1.03(0.98,1.09) | 1.00(0.94,1.05) | 1.00(0.94,1.05) |
|  | R2 | 1.79e-5 | 0.0009 | 0.0003 | 0.0004 | 5.33e-6 | 1.40e-6 |
|  | p | 0.795 | 0.0694 | 0.266 | 0.241 | 0.887 | 0.942 |

dep = depression, irb = irritability, psy = psychosis, apt = apathy, VAB = violent/aggressive behaviour. POB= perseverative/obsessive behaviour. cog=cognitive impairment.

**Supplementary Table 7:** All PRS-symptom analyses for OCD PRS

| Symptom | Statistic | PRS cutoff | | | | | |
| --- | --- | --- | --- | --- | --- | --- | --- |
|  |  | 0.0001 | 0.001 | 0.01 | 0.05 | 0.5 | 1 |
| dep | OR | 1.04(0.98,1.10) | 1.00(0.95,1.06) | 1.09(1.03,1.15) | 1.07(1.01,1.14) | 1.08(1.01,1.14) | 1.08(1.02,1.14) |
|  | R2 | 0.0005 | 6.39e-6 | 0.0021 | 0.0016 | 0.0016 | 0.0017 |
|  | p | 0.170 | 0.878 | 0.00516 | 0.0157 | 0.0142 | 0.0129 |
| Irb | OR | 1.02(0.96,1.08) | 1.03(0.98,1.09) | 1.02(0.96,1.08) | 1.01(0.96,1.07) | 1.03(0.97,1.09) | 1.02(0.97,1.08) |
|  | R2 | 9.30e-5 | 0.0004 | 0.0001 | 5.42e-5 | 0.0002 | 0.0002 |
|  | p | 0.556 | 0.237 | 0.478 | 0.653 | 0.380 | 0.416 |
| psy | OR | 1.07(0.98,1.16) | 0.98(0.90,1.07) | 1.05(0.96,1.15) | 0.99(0.91,1.08) | 1.00(0.91,1.09) | 1.00(0.91,1.09) |
|  | R2 | 0.0009 | 7.44e-5 | 0.0005 | 2.50e-5 | 5.20e-6 | 1.09e-6 |
|  | p | 0.137 | 0.665 | 0.284 | 0.802 | 0.909 | 0.958 |
| apt | OR | 1.01(0.96,1.07) | 1.03(0.98,1.09) | 0.99(0.94,1.05) | 0.99(0.93,1.04) | 1.00(0.95,1.06) | 1.00(0.94,1.06) |
|  | R2 | 7.20e-5 | 0.0004 | 1.13e-5 | 6.54e-5 | 1.60e-7 | 8.52e-7 |
|  | p | 0.603 | 0.251 | 0.837 | 0.620 | 0.980 | 0.955 |
| VAB | OR | 1.02(0.96,1.08) | 1.01(0.95,1.07) | 1.03(0.97,1.10) | 1.01(0.95,1.07) | 1.01(0.95,1.07) | 1.01(0.95,1.07) |
|  | R2 | 0.0001 | 3.73e-5 | 0.0003 | 3.79e-5 | 2.50e-5 | 2.00e-5 |
|  | p | 0.498 | 0.716 | 0.298 | 0.714 | 0.766 | 0.790 |
| POB | OR | 1.02(0.96,1.08) | 1.07(1.01,1.13) | 1.06(1.00,1.12) | 1.05(0.99,1.11) | 1.07(1.01,1.14) | 1.07(1.01,1.13) |
|  | R2 | 0.0001 | 0.0015 | 0.0009 | 0.0006 | 0.0015 | 0.0013 |
|  | p | 0.515 | 0.0190 | 0.0667 | 0.126 | 0.0194 | 0.0281 |
| cog | OR | 1.00(0.95,1.06) | 1.03(0.97,1.09) | 1.01(0.96,1.07) | 1.00(0.95,1.06) | 1.00(0.95,1.06) | 1.00(0.95,1.06) |
|  | R2 | 4.78e-9 | 0.0003 | 4.94e-5 | 5.80e-6 | 7.45e-6 | 4.51e-6 |
|  | p | 0.997 | 0.298 | 0.666 | 0.882 | 0.867 | 0.896 |

dep = depression, irb = irritability, psy = psychosis, apt = apathy, VAB = violent/aggressive behaviour. POB= perseverative/obsessive behaviour. cog=cognitive impairment.

**Supplementary Table 8:** All PRS-symptom analyses for AD PRS

| Symptom | Statistic | PRS cutoff | | | | | |
| --- | --- | --- | --- | --- | --- | --- | --- |
|  |  | 0.0001 | 0.001 | 0.01 | 0.05 | 0.5 | 1 |
| dep | OR | 0.97(0.92,1.03) | 0.98(0.92,1.03) | 0.95(0.91,1.02) | 0.98(0.93,1.04) | 1.01(0.96,1.07) | 1.02(0.96,1.08) |
|  | R2 | 0.0003 | 0.0002 | 0.0005 | 8.41e-5 | 4.77e-5 | 0.0001 |
|  | p | 0.303 | 0.427 | 0.171 | 0.579 | 0.676 | 0.490 |
| irb | OR | 1.00(0.95,1.06) | 1.01(0.95,1.06) | 0.99(0.93,1.04) | 1.00(0.94,1.05) | 1.03(0.97,1.09) | 1.03(0.97,1.09) |
|  | R2 | 7.63e-6 | 1.00e-5 | 7.27e-5 | 5.57e-6 | 0.0003 | 0.0002 |
|  | p | 0.866 | 0.847 | 0.602 | 0.885 | 0.316 | 0.346 |
| psy | OR | 1.05(0.97,1.15) | 1.10(1.01,1.20) | 1.08(0.99,1.18) | 1.06(0.97,1.16) | 1.04(0.96,1.14) | 1.04(0.96,1.14) |
|  | R2 | 0.0006 | 0.0019 | 0.0013 | 0.0007 | 0.0003 | 0.0004 |
|  | p | 0.234 | 0.0265 | 0.0729 | 0.181 | 0.350 | 0.326 |
| apt | OR | 1.00(0.94,1.05) | 0.97(0.92,1.03) | 0.99(0.93,1.04) | 1.01(0.95,1.07) | 1.03(0.98,1.09) | 1.03(0.98,1.09) |
|  | R2 | 5.56e-6 | 0.0003 | 6.26e-5 | 2.35e-5 | 0.0003 | 0.0003 |
|  | p | 0.885 | 0.290 | 0.627 | 0.766 | 0.272 | 0.279 |
| VAB | OR | 1.00(0.94,1.06) | 1.00(0.94,1.06) | 1.01(0.95,1.07) | 1.03(0.97,1.09) | 1.02(0.97,1.09) | 1.03(0.97,1.09) |
|  | R2 | 3.17e-6 | 7.47e-6 | 1.56e-5 | 0.0003 | 0.0002 | 0.0002 |
|  | p | 0.916 | 0.871 | 0.814 | 0.316 | 0.429 | 0.393 |
| POB | OR | 1.00(0.95,1.06) | 1.04(0.98,1.10) | 1.05(0.99,1.11) | 1.05(0.99,1.11) | 1.05(0.99,1.11) | 1.04(0.99,1.11) |
|  | R2 | 1.73e-6 | 0.0005 | 0.0007 | 0.0007 | 0.0008 | 0.0006 |
|  | p | 0.937 | 0.165 | 0.114 | 0.119 | 0.0922 | 0.143 |
| cog | OR | 1.00(0.95,1.06) | 1.02(0.96,1.08) | 0.99(0.93,1.04) | 1.00(0.94,1.05) | 1.00(0.95,1.06) | 1.01(0.95,1.07) |
|  | R2 | 1.31e-7 | 0.0001 | 5.00e-5 | 8.62e-6 | 1.68e-6 | 2.89e-5 |
|  | p | 0.982 | 0.503 | 0.664 | 0.857 | 0.937 | 0.741 |

dep = depression, irb = irritability, psy = psychosis, apt = apathy, VAB = violent/aggressive behaviour. POB= perseverative/obsessive behaviour. cog=cognitive impairment.

**Supplementary Table 9:** All PRS-symptom analyses for PD PRS

| Symptom | Statistic | PRS cutoff | | | | | |
| --- | --- | --- | --- | --- | --- | --- | --- |
|  |  | 0.0001 | 0.001 | 0.01 | 0.05 | 0.5 | 1 |
| dep | OR | 0.99(0.94,1.05) | 0.95(0.89,1.00) | 0.96(0.91,1.02) | 0.97(0.91,1.03) | 1.00(0.94,1.06) | 1.00(0.94,1.06) |
|  | R2 | 1.60e-5 | 0.0009 | 0.0005 | 0.0003 | 4.90e-6 | 3.87e-6 |
|  | P | 0.809 | 0.0620 | 0.169 | 0.287 | 0.893 | 0.905 |
| irb | OR | 1.02(0.97,1.08) | 0.99(0.93,1.04) | 0.99(0.94,1.05) | 0.97(0.92,1.03) | 0.98(0.93,1.04) | 0.98(0.93,1.04) |
|  | R2 | 0.0002 | 5.73e-5 | 2.24e-5 | 0.0002 | 0.0001 | 9.03e-5 |
|  | p | 0.403 | 0.643 | 0.772 | 0.338 | 0.521 | 0.561 |
| psy | OR | 1.01(0.92,1.10) | 0.94(0.87,1.03) | 1.01(0.92,1.10) | 1.01(0.93,1.10) | 1.08(0.99,1.18) | 1.08(0.99,1.18) |
|  | R2 | 1.15e-5 | 0.0006 | 1.51e-5 | 1.78e-5 | 0.0011 | 0.0011 |
|  | p | 0.865 | 0.203 | 0.845 | 0.832 | 0.0909 | 0.0960 |
| apt | OR | 1.01(0.96,1.07) | 0.96(0.90,1.01) | 0.99(0.93,1.04) | 1.01(0.95,1.06) | 0.99(0.93,1.04) | 0.98(0.93,1.04) |
|  | R2 | 2.83e-5 | 0.0007 | 6.36e-5 | 9.47e-6 | 5.47e-5 | 8.16e-5 |
|  | p | 0.744 | 0.108 | 0.625 | 0.850 | 0.650 | 0.580 |
| VAB | OR | 1.05(0.99,1.12) | 1.00(0.94,1.06) | 0.99(0.93,1.05) | 0.98(0.92,1.04) | 0.98(0.93,1.04) | 0.98(0.93,1.05) |
|  | R2 | 0.0009 | 2.39e-6 | 5.05e-5 | 0.0002 | 8.71e-5 | 7.62e-5 |
|  | p | 0.0798 | 0.927 | 0.672 | 0.419 | 0.578 | 0.603 |
| POB | OR | 1.03(0.98,1.09) | 0.96(0.91,1.02) | 1.00(0.95,1.06) | 0.97(0.92,1.03) | 0.98(0.93,1.04) | 0.98(0.92,1.04) |
|  | R2 | 0.0004 | 0.0006 | 7.50e-6 | 0.0002 | 0.0001 | 0.0002 |
|  | p | 0.246 | 0.155 | 0.868 | 0.366 | 0.507 | 0.446 |
| cog | OR | 1.09(1.03,1.16) | 1.02(0.96,1.08) | 1.04(0.99,1.10) | 1.01(0.96,1.07) | 0.99(0.94,1.05) | 0.99(0.93,1.04) |
|  | R2 | 0.0026 | 0.0001 | 0.0006 | 5.40e-5 | 1.78e-5 | 5.59e-5 |
|  | p | 0.00167 | 0.537 | 0.137 | 0.652 | 0.795 | 0.646 |

dep = depression, irb = irritability, psy = psychosis, apt = apathy, VAB = violent/aggressive behaviour. POB= perseverative/obsessive behaviour. cog=cognitive impairment.

**Supplementary Table 10:** All PRS-symptom analyses for intelligence PRS

| Symptom | Statistic | PRS cutoff | | | | | |
| --- | --- | --- | --- | --- | --- | --- | --- |
|  |  | 0.0001 | 0.001 | 0.01 | 0.05 | 0.5 | 1 |
| dep | OR | 0.95(0.90,1.01) | 0.93(0.87,0.98) | 0.95(0.89,1.00) | 0.94(0.89,1.00) | 0.93(0.88,0.99) | 0.93(0.88,0.99) |
|  | R2 | 0.0008 | 0.0018 | 0.0010 | 0.0011 | 0.0016 | 0.0016 |
|  | P | 0.0799 | 0.00951 | 0.0593 | 0.0410 | 0.0152 | 0.0166 |
| irb | OR | 0.92(0.87,0.97) | 0.93(0.88,0.99) | 0.91(0.86,0.96) | 0.90(0.85,0.95) | 0.90(0.85,0.95) | 0.90(0.85,0.95) |
|  | R2 | 0.0024 | 0.0016 | 0.0032 | 0.0034 | 0.0036 | 0.0037 |
|  | p | 0.00306 | 0.0139 | 0.000532 | 0.000344 | 0.000246 | 0.000199 |
| psy | OR | 0.90(0.82,0.98) | 0.93(0.85,1.01) | 0.94(0.86,1.02) | 0.95(0.87,1.04) | 0.91(0.84,1.00) | 0.91(0.84,0.99) |
|  | R2 | 0.0024 | 0.0010 | 0.0008 | 0.0004 | 0.0017 | 0.0017 |
|  | p | 0.0147 | 0.104 | 0.159 | 0.295 | 0.0405 | 0.0371 |
| apt | OR | 0.94(0.89,1.00) | 0.94(0.88,0.99) | 0.93(0.88,0.98) | 0.91(0.86,0.96) | 0.90(0.85,0.95) | 0.90(0.85,0.95) |
|  | R2 | 0.0012 | 0.0015 | 0.0020 | 0.0030 | 0.0039 | 0.0041 |
|  | p | 0.0341 | 0.0166 | 0.00650 | 0.000852 | 0.000118 | 9.27e-5 |
| VAB | OR | 0.91(0.86,0.97) | 0.90(0.85,0.96) | 0.88(0.83,0.93) | 0.89(0.84,0.95) | 0.88(0.83,0.93) | 0.88(0.83,0.93) |
|  | R2 | 0.0027 | 0.0031 | 0.0050 | 0.0040 | 0.0048 | 0.0048 |
|  | p | 0.00198 | 0.00101 | 2.44e-5 | 0.000156 | 4.09e-5 | 3.94e-5 |
| POB | OR | 0.96(0.91,1.02) | 0.96(0.91,1.02) | 0.93(0.88,0.98) | 0.93(0.88,0.99) | 0.94(0.89,1.00) | 0.94(0.88,0.99) |
|  | R2 | 0.0006 | 0.0006 | 0.0018 | 0.0016 | 0.0013 | 0.0014 |
|  | p | 0.150 | 0.152 | 0.0103 | 0.0149 | 0.0318 | 0.0247 |
| cog | OR | 0.92(0.87,0.97) | 0.91(0.86,0.96) | 0.89(0.84,0.94) | 0.90(0.85,0.95) | 0.90(0.85,0.96) | 0.90(0.85,0.96) |
|  | R2 | 0.0025 | 0.0031 | 0.0045 | 0.0040 | 0.0034 | 0.0033 |
|  | p | 0.00213 | 0.000633 | 3.90e-5 | 0.000108 | 0.000378 | 0.000384 |

dep = depression, irb = irritability, psy = psychosis, apt = apathy, VAB = violent/aggressive behaviour. POB= perseverative/obsessive behaviour. cog=cognitive impairment.

**Supplementary Table 11**

Testing for sex differences in PRS odds ratios for symptoms with both a significant sex difference and a significant PRS association

| Symptom | PRS | cutoff | All | | Males | | Females | | M vs F  P value |
| --- | --- | --- | --- | --- | --- | --- | --- | --- | --- |
|  |  |  | OR | p | OR | p | OR | P |  |
| depression | SCZ | 0.05 | 1.12 | 2.47e-4 | 1.16 | 3.79e-4 | 1.08 | 0.0894 | 0.207 |
|  | BPD | 0.01 | 1.11 | 3.06e-4 | 1.17 | 1.96e-4 | 1.06 | 0.180 | 0.114 |
|  | MDD | 0.01 | 1.17 | 9.01e-8 | 1.18 | 4.14e-5 | 1.13 | 3.51e-3 | 0.386 |
|  | INT | 0.001 | 0.93 | 9.51e-3 | 0.92 | 0.0342 | 0.94 | 0.135 | 0.719 |
| irritability | SCZ | 0.05 | 1.17 | 1.03e-7 | 1.15 | 1.44e-3 | 1.18 | 2.74e-5 | 0.598 |
|  | MDD | 0.001 | 1.12 | 4.42e-5 | 1.13 | 4.19e-3 | 1.12 | 6.18e-3 | 0.869 |
|  | INT | 1 | 0.90 | 1.99e-4 | 0.93 | 0.0966 | 0.87 | 3.42e-4 | 0.202 |
| VAB | SCZ | 0.05 | 1.13 | 5.46e-5 | 1.14 | 3.49e-3 | 1.13 | 6.59e-3 | 0.904 |
|  | BPD | 0.001 | 1.11 | 5.35e-4 | 1.07 | 0.0961 | 1.16 | 7.73e-4 | 0.203 |
|  | ADHD | 0.05 | 1.09 | 5.26e-3 | 1.08 | 0.0711 | 1.12 | 0.0131 | 0.588 |
|  | MDD | 0.001 | 1.13 | 8.45e-5 | 1.10 | 0.0243 | 1.15 | 1.41e-3 | 0.436 |
|  | INT | 0.01 | 0.88 | 2.44e-5 | 0.88 | 2.22e-3 | 0.88 | 4.08e-3 | 0.999 |

VAB = violent/aggressive behaviour.

**Supplementary Table 12:** Logistic regression of symptom on multiple PRS simultaneously, correcting for the other symptoms

| SYMPTOM | PRS | PRS CUT-OFF | OR | P VALUE |
| --- | --- | --- | --- | --- |
| Depression | **MDD**  Schizophrenia  BPD  Intelligence | **0.01**  0.05  0.01  0.001 | **1.14**  1.04  1.07  0.99 | **8.52x10^-5^**  0.253  0.0617  0.679 |
| Irritability | **Schizophrenia**  MDD  Intelligence | **0.05**  0.001  1 | **1.09**  1.07  0.97 | **0.0164**  0.117  0.384 |
| Psychosis | **Schizophrenia**  MDD  **Alzheimer’s**  Intelligence | **0.001**  0.05  **0.001**  0.0001 | **1.14**  1.05  **1.13**  0.95 | **0.00688**  0.302  **0.0139**  0.296 |
| VAB | Intelligence | 0.01 | 0.93 | 0.0630 |
|  | Schizophrenia | 0.05 | 1.03 | 0.480 |
|  | MDD | 0.001 | 1.04 | 0.272 |
|  | **BPD** | **0.001** | **1.14** | **9.11x10^-4^** |
|  | ADHD | 0.05 | 1.05 | 0.209 |
|  | ASD | 0.01 | 1.04 | 0.262 |
| POB | Schizophrenia  BPD  Intelligence  MDD  OCD | 0.01  0.5  0.01  0.5  0.001 | 1.03 | 0.446 |
|  |  |  | 1.06 | 0.0725 |
|  |  |  | 0.97 | 0.393 |
|  |  |  | 1.01 | 0.783 |
|  |  |  | 1.07 | 0.0532 |
| Apathy | Intelligence  MDD  BPD  Schizophrenia  ASD | 1  0.05  0.001  0.0001  0.001 | 0.95  1.00  1.05  1.03  1.06 | 0.137  0.980  0.110  0.380  0.0808 |
| Cognitive | **Intelligence** | **0.01** | **0.91** | **0.00411** |
| impairment | **Parkinson's** | **0.0001** | **1.10** | **0.00355** |
|  | MDD | 0.5 | 1.03 | 0.319 |

Symptom-PRS pairs included in the analysis if they reach nominal significance (p<0.05) when analysed alone. The PRS cutoff giving the most significant association was used. Symptom-PRS pairs shown in **bold** reach nominal significance (p<0.05) after correction both for the other PRS shown in the table and for the other symptoms.

VAB= violent/aggressive behaviour, POB=perserverative/obsessive behaviour. MDD= major depressive disorder. BPD= bipolar disorder. ASD= autism spectrum disorder. ADHD = attention deficit hyperactivity disorder. OR= Odds ratio for presence of symptom associated with 1 s.d. increase in PRS.

**Supplementary Table 13** Association of PRS with number of symptoms

| PRS | Number of symptoms | | | | | | | | P(fac) | P(quant) |
| --- | --- | --- | --- | --- | --- | --- | --- | --- | --- | --- |
|  | 0 (473) | 1  (700) | 2  (858) | 3  (952) | 4  (865) | 5  (715) | 6  (433) | 7  (164) |  |  |
| SCZ-0001 | -0.090 | -0.076 | -0.024 | 0.002 | 0.051 | -0.010 | 0.142 | 0.118 | 0.00285 | 4.80e-5 |
| SCZ-001 | -0.091 | -0.076 | -0.027 | -0.002 | 0.039 | 0.024 | 0.162 | 0.109 | 0.00134 | 6.48e-6 |
| SCZ-01 | -0.114 | -0.094 | -0.064 | 0.024 | 0.018 | 0.083 | 0.167 | 0.138 | 5.62e-6 | 5.06e-9 |
| SCZ-05 | -0.128 | -0.076 | -0.068 | 0.023 | 0.026 | 0.069 | 0.164 | 0.166 | 5.88e-6 | 4.45e-9 |
| SCZ-5 | -0.108 | -0.068 | -0.064 | 0.036 | 0.003 | 0.070 | 0.132 | 0.206 | 4.62e-5 | 1.11e-7 |
| SCZ-1 | -0.112 | -0.069 | -0.059 | 0.032 | 0.005 | 0.069 | 0.128 | 0.024 | 6.67e-5 | 1.25e-7 |
| BIP-0001 | -0.039 | -0.069 | 0.001 | -0.004 | 0.047 | -0.014 | 0.065 | -0.021 | 0.327 | 0.067 |
| BIP-001 | -0.071 | -0.066 | -0.024 | 0.024 | 0.042 | 0.005 | 0.090 | 0.026 | 0.109 | 0.00345 |
| BIP-01 | -0.091 | -0.053 | -0.020 | 0.039 | 0.012 | 0.024 | 0.092 | 0.032 | 0.102 | 0.00268 |
| BIP-05 | -0.086 | -0.035 | -0.002 | 0.028 | -0.011 | 0.016 | 0.069 | 0.101 | 0.250 | 0.0118 |
| BIP-5 | -0.067 | -0.030 | 0.008 | -0.005 | -0.032 | 0.034 | 0.047 | 0.107 | 0.396 | 0.0348 |
| BIP-1 | -0.061 | -0.029 | 0.0002 | -0.011 | -0.033 | 0.040 | 0.044 | 0.120 | 0.354 | 0.0273 |
| MDD-0001 | -0.127 | -0.082 | -0.003 | 0.032 | -0.010 | 0.067 | -0.006 | 0.102 | 0.0122 | 0.000897 |
| MDD-001 | -0.178 | -0.069 | -0.014 | 0.052 | -0.008 | 0.081 | 0.031 | 0.192 | 4.32e-5 | 2.583e-6 |
| MDD-01 | -0.167 | -0.074 | -0.015 | -0.004 | 0.034 | 0.059 | 0.053 | 0.261 | 2.46e-5 | 1.34e-7 |
| MDD-05 | -0.175 | -0.082 | -0.023 | -0.014 | 0.013 | 0.067 | 0.084 | 0.206 | 3.25e-5 | 4.58e-8 |
| MDD-5 | -0.133 | -0.117 | -0.011 | -0.013 | 0.023 | 0.058 | 0.077 | 0.263 | 1.34e-5 | 4.17e-8 |
| MDD-1 | -0.136 | -0.116 | -0.011 | -0.011 | 0.024 | 0.057 | 0.072 | 0.253 | 2.15e-5 | 6.35e-8 |
| ADHD-0001 | -0.030 | 0.016 | 0.012 | -0.002 | 0.022 | -0.034 | -0.022 | 0.020 | 0.945 | 0.800 |
| ADHD-001 | -0.024 | 0.000 | 0.005 | -0.024 | 0.034 | 0.004 | -0.033 | 0.076 | 0.867 | 0.602 |
| ADHD-01 | -0.059 | -0.039 | 0.006 | 0.008 | 0.030 | 0.017 | 0.015 | -0.046 | 0.762 | 0.203 |
| ADHD-05 | -0.061 | -0.043 | 0.008 | -0.020 | 0.039 | 0.002 | 0.038 | 0.049 | 0.580 | 0.0561 |
| ADHD-5 | -0.056 | -0.032 | -0.014 | -0.034 | 0.079 | 0.002 | 0.040 | 0.020 | 0.235 | 0.0464 |
| ADHD-1 | -0.055 | -0.029 | -0.009 | -0.035 | 0.083 | -0.002 | 0.028 | 0.009 | 0.238 | 0.0821 |
| ASD-0001 | -0.026 | 0.089 | -0.040 | -0.010 | -0.014 | -0.001 | -0.069 | 0.123 | 0.104 | 0.586 |
| ASD-001 | -0.004 | 0.040 | -0.041 | -0.043 | -0.021 | 0.079 | 0.052 | 0.120 | 0.101 | 0.102 |
| ASD-01 | -0.015 | 0.024 | -0.045 | -0.022 | -0.013 | 0.055 | 0.048 | -0.025 | 0.547 | 0.301 |
| ASD-05 | -0.002 | 0.015 | -0.024 | -0.074 | -0.004 | 0.045 | 0.025 | 0.004 | 0.426 | 0.470 |
| ASD-5 | 0.041 | 0.010 | -0.034 | -0.076 | -0.013 | 0.010 | 0.039 | -0.026 | 0.397 | 0.988 |
| ASD-1 | 0.044 | 0.008 | -0.034 | -0.075 | -0.017 | 0.008 | 0.038 | -0.025 | 0.409 | 0.958 |
| OCD-0001 | -0.066 | 0.020 | -0.006 | 0.017 | 0.010 | -0.011 | 0.009 | 0.152 | 0.490 | 0.209 |
| OCD-001 | -0.046 | -0.001 | 0.001 | 0.005 | 0.020 | -0.007 | 0.053 | 0.073 | 0.856 | 0.169 |
| OCD-01 | -0.031 | -0.041 | 0.023 | -0.010 | 0.050 | 0.009 | 0.071 | 0.005 | 0.523 | 0.0895 |
| OCD-05 | -0.035 | -0.017 | 0.025 | 0.007 | 0.023 | 0.017 | 0.053 | -0.092 | 0.740 | 0.453 |
| OCD-5 | -0.038 | -0.021 | 0.021 | 0.011 | -0.001 | 0.026 | 0.130 | -0.101 | 0.159 | 0.137 |
| OCD-1 | -0.033 | -0.025 | 0.022 | 0.006 | -0.002 | 0.028 | 0.126 | -0.107 | 0.169 | 0.157 |
| INT-0001 | 0.114 | 0.101 | -0.033 | 0.057 | -0.038 | -0.025 | -0.038 | -0.230 | 2.17e-4 | 4.15e-5 |
| INT-001 | 0.100 | 0.112 | -0.020 | 0.025 | -0.025 | -0.030 | -0.086 | -0.161 | 1.88e-3 | 2.81e-5 |
| INT-01 | 0.121 | 0.118 | -0.006 | 0.030 | -0.038 | -0.054 | -0.113 | -0.120 | 1.61e-4 | 6.25e-7 |
| INT-05 | 0.123 | 0.116 | -0.003 | 0.078 | -0.074 | -0.031 | -0.136 | -0.072 | 7.19e-6 | 5.26e-7 |
| INT-5 | 0.093 | 0.113 | 0.014 | 0.059 | -0.090 | -0.027 | -0.156 | -0.103 | 1.24e-6 | 7.46e-8 |
| INT-1 | 0.092 | 0.137 | 0.016 | 0.062 | -0.089 | -0.029 | -0.160 | -0.097 | 7.93e-7 | 5.78e-8 |

Number of symptoms analysed both as an 8-level factor, denoted by p(fac) and as a quantitative variable, denoted by p(quant). Numbers in columns headed by symptom count are the mean PRS among individuals with that number of symptoms.

**Supplementary Table 14**.

Association of SCZ PRS with symptoms after correction for the number of other symptoms present (general severity)

| PRS |  |  |  |  |  |  |  |
| --- | --- | --- | --- | --- | --- | --- | --- |
|  | psy | dep | irb | VAB | apt | POB | cog |
| SCZ-0001 | **0.00416** | 0.234 | 0.138 | 0.914 | 0.458 | 0.187 | 0.253 |
| SCZ-001 | **0.00370** | 0.158 | 0.0662 | 0.717 | 0.850 | **0.0207** | 0.470 |
| SCZ-01 | **0.0157** | 0.104 | **0.00293** | 0.227 | 0.548 | 0.0589 | 0.929 |
| SCZ-05 | **0.0383** | 0.0426 | **0.000904** | 0.0644 | 0.339 | 0.269 | 0.975 |
| SCZ-5 | **0.0249** | 0.0844 | **0.00370** | 0.189 | 0.204 | **0.0432** | 0.694 |
| SCZ-1 | **0.0295** | 0.0824 | **0.00419** | 0.176 | 0.195 | **0.0412** | 0.733 |

Nominally significant associations shown in **bold**.

dep = depression, irb = irritability, psy = psychosis, apt = apathy, VAB = violent/aggressive behaviour. POB= perseverative/obsessive behaviour. cog=cognitive impairment.

**Supplementary Table 15**.

Association of MDD PRS with symptoms after correction for the number of other symptoms present (general severity)

| PRS |  |  |  |  |  |  |  |
| --- | --- | --- | --- | --- | --- | --- | --- |
|  | psy | dep | irb | VAB | apt | POB | cog |
| MDD-0001 | 0.350 | **0.00369** | **0.00298** | 0.0667 | 0.308 | 0.193 | 0.600 |
| MDD-001 | 0.113 | **0.00306** | **0.0183** | **0.0234** | 0.468 | 0.243 | 0.318 |
| MDD-01 | 0.152 | **2.41x10^-5^** | 0.0589 | 0.0588 | 0.885 | 0.602 | 0.945 |
| MDD-05 | 0.115 | **3.46x10^-4^** | 0.196 | 0.258 | 0.759 | 0.749 | 0.202 |
| MDD-5 | 0.234 | **9.34x10^-4^** | 0.388 | 0.101 | 0.909 | 0.783 | 0.195 |
| MDD-1 | 0.290 | **5.44x10^-4^** | 0.336 | 0.0906 | 0.863 | 0.898 | 0.246 |

Nominally significant associations shown in **bold**.

dep = depression, irb = irritability, psy = psychosis, apt = apathy, VAB = violent/aggressive behaviour. POB= perseverative/obsessive behaviour. cog=cognitive impairment.

**Supplementary Table 16.**

Association of intelligence PRS with symptoms after correction for the number of other symptoms present (general severity)

| PRS |  |  |  |  |  |  |  |
| --- | --- | --- | --- | --- | --- | --- | --- |
|  | psy | dep | irb | VAB | apt | POB | cog |
| INT-0001 | 0.179 | 0.862 | 0.277 | 0.110 | 0.544 | 0.842 | **0.036** |
| INT-001 | 0.637 | 0.221 | 0.701 | 0.088 | 0.429 | 0.729 | **0.022** |
| INT-01 | 0.988 | 0.848 | 0.167 | **0.019** | 0.570 | 0.551 | **0.0078** |
| INT-05 | 0.696 | 0.728 | 0.149 | 0.069 | 0.149 | 0.654 | **0.0113** |
| INT-5 | 0.553 | 0.448 | 0.166 | **0.044** | 0.069 | 0.939 | **0.0444** |
| INT-1 | 0.535 | 0.487 | 0.149 | **0.047** | 0.065 | 0.981 | **0.0476** |

Nominally significant associations shown in **bold**.

dep = depression, irb = irritability, psy = psychosis, apt = apathy, VAB = violent/aggressive behaviour. POB= perseverative/obsessive behaviour. cog=cognitive impairment.

**Appendix A**

**Guidelines for the age at onset questionnaire for EHDN-HD Registry patients**

This questionnaire will capture essential and detailed information on when signs and symptoms that may be related or unrelated to HD first appear. Use the suggested wording below as a guide, but ask any other questions you need to try and date the onset as accurately as possible. Ask the patient first, then the carer (explain to the carer at the beginning that you will give them a chance to state their views after you have first heard from the patient) and finally record your own judgement about onset, taking account of what has been said by both patient and carer, and also what you can find in the case notes (or any other independent sources of information). It is not uncommon to find a record by a genetic counsellor who saw the patient and noticed choreiform movements, or an interview with a relative who mentioned that the family think the patient is affected, and this information may pre-date the estimates of either the patient or the carer who is acting as your informant on this occasion. Use your own judgement about the evidence, but avoid making guesses based only on your own experience of how long the average patient would need to be affected to reach this stage in the illness.

1. Motor symptoms

When did you first notice any motor symptoms of HD, such as fidgety movements or clumsiness? Can you remember a particular occasion when this happened and you first realised you might be developing the family illness? What were you doing at the time? Where were you? Did you mention it to anyone else? (Try to identify any landmark events in the patient’s life around that time, such as holidays, births, marriages & deaths in the family, etc, which might help to anchor the onset date by asking whether the onset was before or after that particular landmark)

2. Depression:

Have you ever suffered from depression for a period of 4 weeks or more? Was there ever a time you felt so low that you thought about ending it all? Have you been treated with antidepressants? When was the first time this happened to you (doesn’t necessarily have to have been medicated in the first or any subsequent instance)?

3. Irritability and aggression:

Have there been times when you were irritable, bad tempered or ‘cranky’ for a period of 4 weeks or more? Have you had difficulty keeping control of your temper? When did this begin? If so, how bad was the worst ever episode? Has there been overt aggression? If so when did this first occur?

4. Apathy:

Have you noticed any change in your level of motivation? For example, has there ever been a period of four weeks or more when you lost interest in things that usually matter to you, or could not be bothered with activities you usually enjoy, or needed pushing to get around to jobs that need to be done, or spent a lot of time sitting around doing nothing? Did this ever happen at a time when you were not also feeling depressed? When did you first notice this change?

5. Perseverative or Obsessive/Compulsive behaviours:

Have you ever been troubled by recurrent thoughts, fears or mental images that came back over and over again, or had urges to do something over and over (e.g. double-checking, counting things, or washing your hands repeatedly)? Or perhaps found yourself unable to stop thinking about something, so that friends said you were getting it out of proportion, or getting a ‘bee in your bonnet’? If so, how bad was the worst ever episode? When was the first time this happened to you?

6. Presence of psychosis:

Has the patient ever reported symptoms resembling schizophrenia e.g. paranoid thinking, delusions and hearing voices. If so when was the first episode?

7. Cognitive symptoms

Have you ever noticed any problems with your memory, or your ability to concentrate on things? Have you found it more difficult than it used to be to organise things, or perhaps noticed that it takes longer to get things done nowadays? Has anybody commented on this, or maybe criticised your performance at work or when doing chores at home? Can you remember the first time this happened? (probe as above and look for evidence such as job changes etc. which might suggest early cognitive difficulties)

**Appendix B**

**European Huntington’s Disease Network REGISTRY Study Investigators**

**Registry Steering committee:** Anne-Catherine Bachoud-Lévi, Anna Rita Bentivoglio, Ida Biunno, Raphael M Bonelli, Jean-Marc Burgunder, Stephen B Dunnett, Joaquim J Ferreira, Olivia J. Handley, Arvid Heiberg, Torsten Illmann, G Bernhard Landwehrmeyer, Jamie Levey, Maria A. Ramos-Arroyo, Jørgen E Nielsen, Susana Pro Koivisto, Markku Päivärinta, Raymund A.C. Roos, Ana Rojo Sebastián, Sarah J Tabrizi, Wim Vandenberghe, Christine Verellen-Dumoulin, Tereza Uhrova, Jan Wahlström, Jacek Zaremba

**Language coordinators:** Verena Baake, Katrin Barth, Monica Bascuñana Garde, Sabrina Betz, Reineke Bos, Jenny Callaghan, Adrien Come, Leonor Correia Guedes, Daniel Ecker, Ana Maria Finisterra, Ruth Fullam, Mette Gilling, Lena Gustafsson, Olivia J. Handley, Carina Hvalstedt, Christine Held, Kerstin Koppers, Claudia Lamanna, Matilde Laurà, Asunción Martínez Descals, Saül Martinez-Horta, Tiago Mestre, Sara Minster, Daniela Monza, Lisanne Mütze, Martin Oehmen, Michael Orth, Hélène Padieu, Laurent Paterski, Nadia Peppa, Susana Pro Koivisto, Martina Di Renzo, Amandine Rialland, Niini Røren, Pavla Šašinková, Erika Timewell, Jenny Townhill, Patricia Trigo Cubillo, [Wildson Vieira](https://imap.uni-ulm.de/horde/imp/message.php?thismailbox=INBOX&mailbox=%2A%2Asearch_3pty3apzsxkw000sow4444&index=135591) da Silva, Marleen R van Walsem, Carina Whalstedt, Marie-Noelle Witjes-Ané, [Grzegorz Witkowski](https://imap.uni-ulm.de/horde/imp/message.php?thismailbox=INBOX&mailbox=%2A%2Asearch_50vnt49yihwkc4cc048c4w&index=149227) , Abigail Wright, Daniel Zielonka, Eugeniusz Zielonka, Paola Zinzi

**AUSTRIA**

**Graz (Medizinische Universitäts Graz, Psychiatrie):** Raphael M Bonelli, Sabine Lilek, Karen Hecht, Brigitte Herranhof, Anna Holl (formerly Hödl), Hans-Peter Kapfhammer, Michael Koppitz, Markus Magnet, Nicole Müller, Daniela Otti, Annamaria Painold, Karin Reisinger, Monika Scheibl, Helmut Schöggl, Jasmin Ullah

**Innsbruck (Universitätsklinik Innsbruck, Neurologie):** Eva-Maria Braunwarth, Florian Brugger, Lisa Buratti, Eva-Maria Hametner, Caroline Hepperger, Christiane Holas, Anna Hotter, Anna Hussl, Christoph Müller, Werner Poewe, Klaus Seppi, Fabienne Sprenger, Gregor Wenning

**BELGIUM**

**Bierbeek:** Andrea Boogaerts, Godelinde Calmeyn, Isabelle Delvaux, Dirk Liessens, Nele Somers

**Charleroi (Institut de Pathologie et de Génétique (IPG)):** Michel Dupuit, Cécile Minet, Dominique van Paemel, Pascale Ribaï, Christine Verellen-Dumoulin

**Leuven:** (Universitair Ziekenhuis Gasthuisberg,): Andrea Boogaerts, Wim Vandenberghe, Dimphna van Reijen

**CZECH REPUBLIC**

**Prague (Extrapyramidové centrum, Neurologická klinika, 1. LF UK a VFN):** Jiří Klempíř, Veronika Majerová, Jan Roth, Irena Stárková

**DENMARK**

**Copenhagen (Neurogenetics Clinic, Danish Dementia Research Centre, Rigshospitalet, University of Copenhagen):** Lena E. Hjermind, Oda Jacobsen, Jørgen E Nielsen, Ida Unmack Larsen, Tua Vinther-Jensen

**FINLAND**

**Turku-Suvituuli (Rehabilitation Centre Suvituuli):** Heli Hiivola, Hannele Hyppönen, Kirsti Martikainen, Katri Tuuha

**FRANCE**

**Angers (Centre de référence des maladies neurogénétique- CHU d’Angers):** Philippe Allain, Dominique Bonneau, Marie Bost, Bénédicte Gohier, Marie-Anne Guérid, Audrey Olivier, Adriana Prundean, Clarisse Scherer-Gagou, Christophe Verny

**Bordeaux (Hôpital CHU Pellegrin):** Blandine Babiloni, Sabrina Debruxelles, Charlotte Duché, Cyril Goizet, Laetitia Jameau, Danielle Lafoucrière, Umberto Spampinato

**Lille-Amiens:**

**Lille (CHRU Roger Salengro) :** Rekha Barthélémy, Christelle De Bruycker, Maryline Cabaret, Anne-Sophie Carette, Eric Decorte  Luc Defebvre, Marie Delliaux, Arnaud Delval, Alain Destee, Kathy Dujardin, Marie-Hélène Lemaire, Sylvie Manouvrier, Mireille Peter, Lucie Plomhouse, Bernard Sablonnière, Clémence Simonin, Stéphanie Thibault-Tanchou, Isabelle Vuillaume

**Amiens (CHU Nord) :** Marcellin Bellonet, Hassan Berrissoul, Stéphanie Blin, Françoise Courtin, Cécile Duru, Véronique Fasquel, Olivier Godefroy, Pierre Krystkowiak, Béatrice Mantaux, Martine Roussel, Sandrine Wannepain

**Marseille (Hôpital La Timone) :** Jean-Philippe Azulay, Marie Delfini, Alexandre Eusebio, Frédérique Fluchere, Laura Mundler

**Strasbourg (Hôpital Civil) :** Mathieu Anheim, Celine Julié, Ouhaid Lagha Boukbiza, Nadine Longato, Gabrielle Rudolf, Christine Tranchant, Marie-Agathe Zimmermann

**GERMANY**

**Aachen (Universitätsklinikum Aachen, Neurologische Klinik):** Christoph Michael Kosinski, Eva Milkereit, Daniela Probst, Kathrin Reetz, Christian Sass, Johannes Schiefer, Christiane Schlangen, Cornelius J. Werner

**Berlin (Klinik und Poliklinik für Neurologie - Charité - Universitätsmedizin Berlin):** Harald Gelderblom, Josef Priller, Harald Prüß, Eike Jakob Spruth

**Bochum (Huntington-Zentrum (NRW) Bochum im St. Josef-Hospital):** Gisa Ellrichmann, Lennard Herrmann, Rainer Hoffmann, Barbara Kaminski, Peter Kotz, Christian Prehn, Carsten Saft

**Dinslaken (Reha Zentrum in Dinslaken im Gesundheitszentrums Lang):** Herwig Lange, Robert Maiwald

**Dresden (Universitätsklinikum Carl Gustav Carus an der Technischen Universität Dresden, Klinik und Poliklinik für Neurologie):** Matthias Löhle, Antonia Maass, Simone Schmidt, Cecile Bosredon, Alexander Storch, Annett Wolz, Martin Wolz

**Freiburg (Universitätsklinik Freiburg, Neurologie):** Philipp Capetian, Johann Lambeck, Birgit Zucker

**Hamburg (Universitätsklinikum Hamburg-Eppendorf, Klinik und Poliklinik für Neurologie):** Kai Boelmans, Christos Ganos, Walburgis Heinicke, Ute Hidding, Jan Lewerenz, Alexander Münchau, Michael Orth, Jenny Schmalfeld, Lars Stubbe, Simone Zittel

**Hannover (Neurologische Klinik mit Klinischer Neurophysiologie, Medizinische Hochschule Hannover):** Gabriele Diercks, Dirk Dressler, Heike Gorzolla, Christoph Schrader, Pawel Tacik

**Itzehoe (Schwerpunktpraxis Huntington, Neurologie und Psychiatrie):** Michael Ribbat

**Marburg KPP (Klinik für Psychiatrie und Psychotherapie Marburg-Süd):** Bernhard Longinus

**Marburg Uni (Universität Marburg, Neurologie):** Katrin Bürk, Jens Carsten Möller, Ida Rissling

**München (Huntington-Ambulanz im Neuro-Kopfzentrum - Klinikum rechts der Isar der Neurologischen Klinik und Poliklinik der Technischen Universität München):** Mark Mühlau, Alexander Peinemann, Michael Städtler, Adolf Weindl, Juliane Winkelmann, Cornelia Ziegler

**Münster (Universitätsklinikum Münster, Klinik und Poliklinik für Neurologie):** Natalie Bechtel, Heike Beckmann, Stefan Bohlen, Eva Hölzner, Herwig Lange, Ralf Reilmann, Stefanie Rohm, Silke Rumpf , Sigrun Schepers, Natalia Weber

**Taufkirchen (Isar-Amper-Klinikum - Klinik Taufkirchen (Vils)):** Matthias Dose, Gabriele Leythäuser, Ralf Marquard, Tina Raab, Alexandra Wiedemann

**Ulm (Universitätsklinikum Ulm, Neurologie):** Katrin Barth, Andrea Buck, Julia Connemann, Daniel Ecker, Carolin Geitner, Christine Held, Andrea Kesse, Bernhard Landwehrmeyer, Christina Lang, Jan Lewerenz, Franziska Lezius, Solveig Nepper, Anke Niess, Michael Orth, Ariane Schneider, Daniela Schwenk, Sigurd Süßmuth, Sonja Trautmann, Patrick Weydt

**ITALY**

**Bari Clinica Neurologica - Neurophysiopatology of Pain Unit UNIVERSITA' DI BARI):** Claudia Cormio, Vittorio Sciruicchio, Claudia Serpino, Marina de Tommaso

**Bologna (DIBINEM - Alma Mater Studiorum - Università di Bologna; IRCCS Istituto delle Scienze Neurologiche di Bologna):** Sabina Capellari, Pietro Cortelli, Roberto Galassi, Rizzo Giovanni, Roberto Poda, Cesa Scaglione

**Florence (Dipartimento di Scienze Neurologiche e Psichiatriche Universita' degli Studi di Firenze-Azienda Ospedaliera Universitaria Careggi):** Elisabetta Bertini, Elena Ghelli, Andrea Ginestroni, Francesca Massaro, Claudia Mechi, Marco Paganini, Silvia Piacentini, Silvia Pradella, Anna Maria Romoli, Sandro Sorbi

**Genoa (Dipartimento di Neuroscienze, Riabilitazione, Oftalmologia, Genetica e Scienze Materno-Infantili, Università di Genova):** Giovanni Abbruzzese, Monica Bandettini di Poggio, Giovanna Ferrandes, Paola Mandich, Roberta Marchese

**Milan (Fondazione IRCCS Istituto Neurologico Carlo Besta):**
Alberto Albanese, Daniela Di Bella, Anna Castaldo, Stefano Di Donato, Cinzia Gellera, Silvia Genitrini, Caterina Mariotti, Daniela Monza, Lorenzo Nanetti, Dominga Paridi, Paola Soliveri, Chiara Tomasello

**Naples (Dipartimento di Neuroscienze, Scienze Riproduttive e Odontostomatologiche, Università Federico II):** Giuseppe De Michele, Luigi Di Maio, Marco Massarelli, Silvio Peluso, Alessandro Roca, Cinzia Valeria Russo, Elena Salvatore, Pierpaolo Sorrentino

**Pozzilli (IS) (Centro di Neurogenetica e Malattie Rare - IRCCS Neuromed):** Enrico Amico, Mariagrazia Favellato, Annamaria Griguoli, Irene Mazzante, Martina Petrollini, Ferdinando Squitieri and **Rome (Lega Italiana Ricerca Huntington e malattie correlate - onlus / www.LIRH.it):** Barbara D'Alessio, Chiara Esposito

**Rome (Istituto di Farmacologia Traslazionale & Istituto di Scienze e Tecnologie della Cognizione /CNR, Istituto di Neurologia Università Cattolica del Sacro Cuore):** Anna Rita Bentivoglio, Marina Frontali, Arianna Guidubaldi, Tamara Ialongo, Gioia Jacopini, Carla Piano, Silvia Romano, Francesco Soleti, Maria Spadaro, Paola Zinzi

**NETHERLANDS**

**Enschede (Medisch Spectrum Twente):** Monique S.E. van Hout, Marloes E. Verhoeven, Jeroen P.P. van Vugt, A. Marit de Weert

**Groningen (Polikliniek Neurologie**): J.J.W. Bolwijn, M. Dekker, B. Kremer, K.L. Leenders, J.C.H. van Oostrom

**Leiden (Leiden University Medical Centre (LUMC**)): Simon J. A. van den Bogaard, Reineke Bos, Eve M. Dumas, Ellen P. ‘t Hart, Raymund A.C. Roos

**Nijmegen (Universitair Medisch Centrum St. Radboud, Neurology**): Berry Kremer, C.C.P. Verstappen

**NORWAY**

**Oslo University Hospital (Rikshospitalet, Dept. of Medical Genetics and Dept. of Neurology):** Olaf Aaserud, Jan Frich C., Arvid Heiberg, Marleen R. van Walsem, Ragnhild Wehus

**Oslo University Hospital (Ulleval**, **Dept. of Medical Genetics and Dept.of Neurorehabilitation)**: Kathrine Bjørgo, Madeleine Fannemel, Per F. Gørvell, Eirin Lorentzen, Susana Pro Koivisto, Lars Retterstøl, Bodil Stokke

**Trondheim (St. Olavs Hospital):** Inga Bjørnevoll, Sigrid Botne Sando

**POLAND**

**Gdansk (St. Adalbert Hospital, Gdansk, Medical University of Gdansk, Neurological and Psychiatric Nursing Dpt.):** Artur Dziadkiewicz, Malgorzata Nowak, Piotr Robowski, Emilia Sitek, Jaroslaw Slawek, Witold Soltan, Michal Szinwelski

**Katowice (Medical University of Silesia, Katowice):** Magdalena Blaszcyk, Magdalena Boczarska-Jedynak, Ewelina Ciach-Wysocka, Agnieszka Gorzkowska, Barbara Jasinska-Myga, Gabriela Kłodowska–Duda, Gregorz Opala, Daniel Stompel

**Krakow (Krakowska Akademia Neurologii):** Krzysztof Banaszkiewicz, Dorota Boćwińska, Kamila Bojakowska-Jaremek, Małgorzata Dec, Malgorzata Krawczyk, Monika Rudzińska, Elżbieta Szczygieł, Andrzej Szczudlik, Anna Wasielewska, Magdalena Wójcik

**Poznan (Poznan University of Medical Sciences, Poland):** Anna Bryl, Anna Ciesielska, Aneta Klimberg, Jerzy Marcinkowski, Husam Samara, Justyna Sempołowicz, Daniel Zielonka

**Warsaw-MU (Medical University of Warsaw, Neurology):** Anna Gogol (formerly Kalbarczyk), Piotr Janik, Hubert Kwiecinski, Zygmunt Jamrozik

**Warsaw-IPiN (Institute of Psychiatry and Neurology Dep. of Genetics, First Dep. of Neurology):** Jakub Antczak, Katarzyna Jachinska, Wioletta Krysa, Maryla Rakowicz, Przemyslaw Richter, Rafal Rola, Danuta Ryglewicz, Halina Sienkiewicz-Jarosz, Iwona Stępniak, Anna Sułek, Grzegorz Witkowski, Jacek Zaremba, Elzbieta Zdzienicka, Karolina Zieora-Jakutowicz

**PORTUGAL**

**Coimbra (Hospital Universitário de Coimbra):** Cristina Januário, Filipa Júlio

**Lisbon (Clinical Pharmacology Unit, Instituto de Medicina Molecular, Faculty of Medicine, University of Lisbon):** Joaquim J Ferreira, Miguel Coelho, Leonor Correia Guedes, Tiago Mendes, Tiago Mestre, Anabela Valadas

**Porto (Hospital de São João, (Faculdade de Medicina da Universidade do Porto)):** Carlos Andrade, Miguel Gago, Carolina Garrett, Maria Rosália Guerra.

**SPAIN**

**Badajoz (Hospital Infanta Cristina):** Carmen Durán Herrera, Patrocinio Moreno Garcia

**Barcelona-Hospital Mútua de Terrassa:** Miquel Aguilar Barbera, Dolors Badenes Guia, Laura Casas Hernanz , Judit López Catena, Pilar Quiléz Ferrer, Ana Rojo Sebastián, Gemma Tome Carruesco

**Barcelona-Bellvitge (Hospital Universitari de Bellvitge):** Jordi Bas, Núria Busquets, Matilde Calopa

**Barcelona-Merced (Hospital Mare de Deu de La Merced):** Misericordia Floriach Robert, Celia Mareca Viladrich, Jesús Miguel Ruiz Idiago, Antonio Villa Riballo

**Burgos (Servicio de Neurología Hospital General Yagüe):** Esther Cubo, Cecilia Gil Polo, Natividad Mariscal Perez, Jessica Rivadeneyra

**Granada (Hospital Universitario San Cecilio, Neurología**): Francisco Barrero, Blas Morales

**Madrid-Clinico (Hospital Clínico Universitario San Carlos):** María Fenollar, Rocío García-Ramos García, Paloma Ortega, Clara Villanueva

**Madrid RYC (Hospital Ramón y Cajal, Neurología):** Javier Alegre, Mónica Bascuñana, Juan Garcia Caldentey, Marta Fatás Ventura, Guillermo García Ribas, Justo García de Yébenes, José Luis López-Sendón Moreno, Patricia Trigo Cubillo

**Madrid FJD (Madrid-Fundación Jiménez Díaz):** Javier Alegre, Fernando Alonso Frech, Justo García de Yébenes, Pedro J García Ruíz, Asunción Martínez-Descals, Rosa Guerrero, María José Saiz Artiga, Vicenta Sánchez

**Murcia (Hospital Universitario Virgen de la Arrixaca):** María Fuensanta Noguera Perea, Lorenza Fortuna, Salvadora Manzanares, Gema Reinante, María Martirio Antequera Torres, Laura Vivancos Moreau

**Oviedo (Hospital Central de Asturias):** Sonia González González, Luis Menéndez Guisasola, Carlos Salvador, Esther Suaréz San Martín

**Palma de Mallorca (Hospital Universitario Son Espases):** Inés Legarda Ramirez, Aranzazú Gorospe, Mónica Rodriguez Lopera, Penelope Navas Arques, María José Torres Rodríguez, Barbara Vives Pastor

**Pamplona (Complejo Hospitalario de Navarra):** Itziar Gaston, Maria Dolores Martinez-Jaurrieta, Maria A. Ramos-Arroyo

**Sevilla ("Hospital Virgen Macarena"):** Jose Manuel Garcia Moreno, Carolina Mendez Lucena, Fatima Damas Hermoso, Eva Pacheco Cortegana, José Chacón Peña, Luis Redondo

**Sevilla (Hospital Universitario Virgen del Rocío):** Fátima Carrillo, María Teresa Cáceres, Pablo Mir, María José Lama Suarez, Laura Vargas-González

**Valencia (Hospital la Fe):** Maria E. Bosca, Francisco Castera Brugada, Juan Andres Burguera, Anabel Campos Garcia, Carmen Peiró Vilaplana

**SWEDEN**

**Göteborg (Sahlgrenska University Hospital**): Peter Berglund, Radu Constantinescu, Gunnel Fredlund, Ulrika Høsterey-Ugander, Petra Linnsand, Liselotte Neleborn-Lingefjärd, Jan Wahlström, Magnus Wentzel

**Umeå (Umeå University Hospital**): Ghada Loutfi, Carina Olofsson, Eva-Lena Stattin, Laila Westman, Birgitta Wikström

**SWITZERLAND**

**Bern:** Jean-Marc Burgunder, Yanik Stebler **(Swiss HD Zentrum)**, Alain Kaelin, Irene Romero, Michael Schüpbach, Sabine Weber Zaugg **(Zentrum für Bewegungsstörungen, Neurologische Klinik und Poliklinik, Universität Bern)**

**Zürich (Department of Neurology, University Hospital Zürich):** Maria Hauer, Roman Gonzenbach, Hans H. Jung, Violeta Mihaylova, Jens Petersen

**UNITED KINGDOM**

**Aberdeen (NHS Grampian Clinical Genetics Centre & University of Aberdeen):** Roisin Jack, Kirsty Matheson, Zosia Miedzybrodzka, Daniela Rae, Sheila A Simpson, Fiona Summers, Alexandra Ure, Vivien Vaughan

**Birmingham (The Barberry Centre, Dept of Psychiatry**): Shahbana Akhtar, Jenny Crooks, Adrienne Curtis, Jenny de Souza (Keylock), John Piedad, Hugh Rickards, Jan Wright

**Bristol (North Bristol NHS Trust, Southmead hospital):** Elizabeth Coulthard, Louise Gethin, Beverley Hayward, Kasia Sieradzan, Abigail Wright

**Cambridge (Cambridge Centre for Brain Repair, Forvie Site):** Matthew Armstrong, Roger A. Barker, Deidre O’Keefe, Anna Di Pietro, Kate Fisher, Anna Goodman, Susan Hill, Ann Kershaw, Sarah Mason, Nicole Paterson, Lucy Raymond, Rachel Swain, Natalie Valle Guzman

**Cardiff (Schools of Medicine and Biosciences, Cardiff University**): Monica Busse, Cynthia Butcher, Jenny Callaghan, Stephen Dunnett, Catherine Clenaghan, Ruth Fullam, Olivia Handley, Sarah Hunt, Lesley Jones, Una Jones, Hanan Khalil, Sara Minster, Michael Owen, Kathleen Price, Anne Rosser, Jenny Townhill

**Edinburgh (Molecular Medicine Centre, Western General Hospital, Department of Clinical Genetics):** Maureen Edwards, Carrie Ho (Scottish Huntington´s Association), Teresa Hughes (Scottish Huntington´s Association), Marie McGill, Pauline Pearson, Mary Porteous, Paul Smith (Scottish Huntington´s Association)

**Fife (Scottish Huntington's Association Whyteman's Brae Hospital):** Peter Brockie, Jillian Foster, Nicola Johns, Sue McKenzie, Jean Rothery, Gareth Thomas, Shona Yates

**Gloucester (Department of Neurology Gloucestershire Royal Hospital):** Liz Burrows, Carol Chu, Amy Fletcher, Deena Gallantrae, Stephanie Hamer, Alison Harding, Stefan Klöppel, Alison Kraus, Fiona Laver, Monica Lewis, Mandy Longthorpe, Ivana Markova, Ashok Raman, Nicola Robertson, Mark Silva, Aileen Thomson, Sue Wild, Pam Yardumian

**Hull (Castle Hill Hospital):** Carol Chu, Carole Evans, Deena Gallentrae, Stephanie Hamer, Alison Kraus, Ivana Markova, Ashok Raman

**Leeds (Chapel Allerton Hospital, Department of Clinical Genetics):** Leeds (Chapel Allerton Hospital, Clinical Genetics): Carol Chu, Stephanie Hamer, Emma Hobson, Stuart Jamieson, Alison Kraus, Ivana Markova, Ashok Raman, Hannah Musgrave, Liz Rowett, Jean Toscano, Sue Wild, Pam Yardumian

**Leicester (Leicestershire Partnership Trust, Mill Lodge):** Colin Bourne, Jackie Clapton, Carole Clayton, Heather Dipple, Dawn Freire-Patino, Janet Grant, Diana Gross, Caroline Hallam, Julia Middleton, Ann Murch, Catherine Thompson

**Liverpool (Walton Centre for Neurology and Neurosurgery**)**:** Sundus Alusi, Rhys Davies, Kevin Foy, Emily Gerrans, Louise Pate

**London (Guy's Hospital):** Thomasin Andrews, Andrew Dougherty, Charlotte Golding, Fred Kavalier, Hana Laing, Alison Lashwood, Dene Robertson, Deborah Ruddy, Alastair Santhouse, Anna Whaite

**London (The National Hospital for Neurology and Neurosurgery**): Thomasin Andrews, Stefania Bruno, Karen Doherty, Charlotte Golding, Salman Haider, Davina Hensman, Nayana Lahiri, Monica Lewis, Marianne Novak, Aakta Patel, Nicola Robertson, Elisabeth Rosser, Sarah Tabrizi, Rachel Taylor, Thomas Warner, Edward Wild

**Manchester (Genetic Medicine, University of Manchester, Manchester Academic Health Sciences Centre and Central Manchester University Hospitals NHS Foundation Trust):** Natalie Arran, Judith Bek, Jenny Callaghan, David Craufurd, Ruth Fullam, Marianne Hare, Liz Howard, Susan Huson, Liz Johnson, Mary Jones, Helen Murphy, Emma Oughton, Lucy Partington-Jones, Dawn Rogers, Andrea Sollom, Julie Snowden, Cheryl Stopford, Jennifer Thompson, Iris Trender-Gerhard, Nichola Verstraelen (formerly Ritchie), Leann Westmoreland

**Oxford (Oxford University Hospitals NHS Trust, Dept. of Neurosciences, University of Oxford):** Richard Armstrong, Kathryn Dixon, Andrea H Nemeth, Gill Siuda, Ruth Valentine

**Plymouth (Plymouth Huntington Disease Service, Mount Gould Hospital):** David Harrison, Max Hughes, Andrew Parkinson, Beverley Soltysiak

**Sheffield (The Royal Hallamshire Hospital– Sheffield Children’s Hospital):** Oliver Bandmann, Alyson Bradbury, Paul Gill, Helen Fairtlough, Kay Fillingham, Isabella Foustanos, Mbombe Kazoka, Kirsty O’Donovan, Nadia Peppa, Cat Taylor, Katherine Tidswell, Oliver Quarrell

**EHDN’s associate site in Singapore: National Neuroscience Institute Singapore:** Jean-Marc Burgunder, Puay Ngoh Lau, Emmanul Pica, Louis Tan

**Appendix C**

**EHDN Behavioural Modifiers Working Group members**

Aad Tibben, Akshay Nair, Andrea Higgins, Andrea Horta, Arvid Heiberg. Asa Peterson, Asunción Martínez Descals, Bernhard Landwehrmeyer, Carolyn Drazinic, Charlotte Goodwin, Cheryl Stopford, Clare Eddy, Constantine Economides, Daniel P. Van Kammen, David Craufurd, Dawn Freire-Patino, Ashraful Bari, Duncan McLauchlan, Emilia Sitek, Erik van Duijn, Fiona Eccles, George El-Nimr, Hanneke Nijsten, Hugh Rickards, Ida Unmack Larsen, Inga Bjørnevoll, James Osborne, Jan Van den Stock, Jan Wright, Jane Simpson, Jennifer Hoblyn, Jenny Callaghan, Jenny De Souza, Jesús Miguel Ruiz Idiago, Jesus Perez Perez, John Maltby, Josef Priller, Karen Anderson, LaVonne Goodman, Lesley Jones, Maria Dale, Marleen van Walsem, Martin Kucharík, Mary Edmondson, Matthias Dose, Mayke Oosterloo, Nicola Robertson, Noora Ovaska, Olga Klempirova, Oliver Tuscher, Pavel Ressner, Philipp Honrath, Rainer Hoffmann, Rebecca Fuller, Saül Martinez Horta, Tereza Uhrova, Tua Vinther-Jensen, Katie Osborne-Crowley, Natascia De Lucia, Marianna Delussi
